# Bivalent interaction through an intrinsically disordered linker promotes transcription activation complex assembly in Notch signaling

**DOI:** 10.1101/2024.11.09.622674

**Authors:** Cyril Cook, Kristen M. Ramsey, Doug Barrick

## Abstract

The Notch signaling pathway regulates cellular differentiation by activating transcription through an unusual heterotrimeric complex comprising the Notch receptor’s intracellular domain (NICD), the DNA-binding protein CSL, and the coactivator MAML. NICD has two binding sites for CSL, a short motif in the RAM region and an ankyrin domain (ANK), connected by an intrinsically disordered linker and which form a bivalent ternary complex with CSL and MAML. Although bivalency is required for maximal transcription activation, the energetic contributions of bivalency and heterotrimer formation within this essential complex are unknown. To elucidate the energetics of bivalency we first determine the free energy of the CSL-ANK-MAML heterotrimer, using isothermal titration calorimetry and developing an obligate heterotrimer model to analyze the data. By comparing this heterotrimerization reaction with binding reactions involving different regions of RAMANK, we determine the energetic contribution of bivalency to heterotrimer assembly. We show that bivalency through the disordered linker increases the effective concentration of ANK, and that the bivalent interaction enhances occupancy of RAM and ANK at their binding sites on CSL by about three orders of magnitude. By redefining the standard state to a lower, more physiological protein concentration, we reveal the importance of the RAMANK intrinsically disordered linker for assembly of the Notch transcription activation complex. This work provides a framework whereby the energetic contributions of intrinsically disordered linkers to higher-order multivalent assembly may be analyzed.

## Introduction

Notch signaling is a signal transduction pathway that is central to cellular differentiation and homeostasis in metazoans, making major contributions to processes such as embryogenesis, wound healing, hematopoesis, neurogenesis, and stem cell maintenance [6, 39]. As such, mutations in Notch signaling genes are associated with various cancers and diseases [1, 5]. Genes comprising the Notch pathway were identified in genetic screens over a century ago [10]. Activation of Notch signaling is initiated by interaction of the Notch transmembrane receptor with ligands on adjacent cells, which triggers proteolysis of the Notch receptor, releasing the Notch intracellular domain (NICD) into the cytosol [24]. NICD is then imported into the nucleus, where it associates with CSL, a DNA-binding protein, and MAML, a co-activator; formation of this NICD:CSL:MAML ternary complex at promoters of Notch responsive genes activates their transcription [20].

Studies over the past 25 years using molecular genetics, biochemistry, and x-ray crystallography have highlighted several unique features of the Notch ternary complex [21]. Most notably, NICD engages CSL through two distinct regions, the RAM region (specifically a short ΦWΦP sequence motif^1^ on the N-terminus of the RAM region) and the ANK domain, a highly conserved set of seven ankyrin repeats (Figure 1B). The ΦWΦP motif of NICD binds to the *β*-trefoil domain (BTD) of CSL, whereas the ANK domain of NICD binds to the N- and C-terminal domains (NTD, CTD) of CSL. The distance between these two binding sites is around 45 °A. The linker separating the ΦWΦP motif and the ANK domain is disordered both in solution and in the crystal structure [26, 36, 3]. X-ray crystallography shows that MAML binds to an extended interface between the ANK domain of NICD and the NTD and CTD domains of CSL as a long kinked *α*-helix [25, 36]. Biochemical studies show that the ANK domain and MAML region mutually stabilize one another in their interaction with CSL, and bind weakly (if at all) on their own [26].

**Figure 1:**
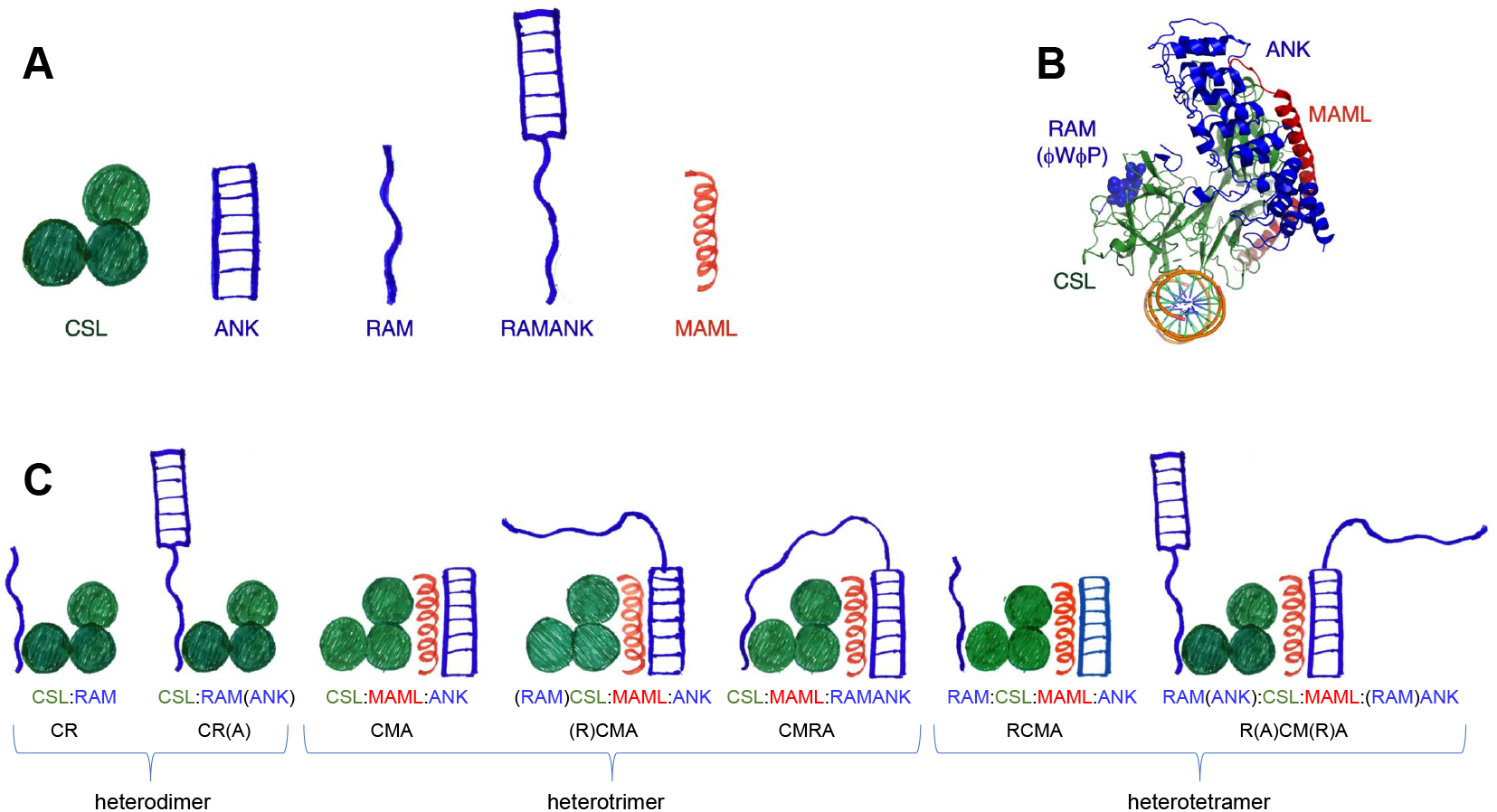
Cartoons of the polypeptides, regions, and domains that form the Notch transcriptional activation complex. (**A**) Individual constructs used in this study. Starting from the left: the three-domain CSL protein in green (residues 25-449), segments from the Notch intracellular domain in blue (ANK, residues 1858-2126; RAM, residues 1758-1888; RAMANK, residues 1758-2126), and the N-terminal helical binding motif of MAML in red (residues 8-76). Boundaries for the constructs used in this study are for human CSL, human Notch1 and human MAML1. (**B**) Crystal structure of the orthologous Notch1 RA-MANK:CSL:MAML1:DNA complex from *C. elegans*, 2fo1.pdb. (**C**) Complexes of CSL with the Notch RAM and ANK region, showing CSL bound both *in cis* and *in trans*, and MAML-bound complexes. The acronyms at the bottom provide a shorthand for the reactions and binding data that follows. The two complexes on the left are dimeric complexes between RAM and CSL. Evidence below and from paralogue studies [28] indicates that the ANK domain does not bind to its site on CSL in the dimeric CSL:RAM(ANK) complex (second from left). In all heterotrimer and heterotetramer complexes, MAML stabilizes ANK binding to CSL. The rightmost heterotrimer (CMRA) displays RAMANK binding to CSL *in cis*, where two separate regions of a single NICD polypeptide engage CSL at two separate sites (comprising a bivalent interaction between RA-MANK and CSL). The two heterotetramer complexes are different versions of both RAM and ANK binding to CSL *in trans*, meaning the RAM and ANK bound regions are from two separate polypeptides.

The relevant segments of these three proteins for ternary complex formation are drawn in Figure 1A, along with the crystal structure of the Notch transcription complex. We refer to the transcription complex formed from the RAMANK region of Notch, CSL, and MAML as “bivalent” because distant regions of NICD simultaneously bind distant sites on CSL. The importance of the bivalent interaction has been demonstrated in transcriptional activation assays in which higher levels of activation are achieved when RAM and ANK are physically connected on a single polypeptide, rather than on two separate polypeptides [17].

Here we determine the energetics of association of Notch RAM, ANK, and RAMANK (from the human Notch1 receptor) with MAML and CSL using isothermal titration calorimetry (ITC). The complexes that we have probed are shown in Figure 1C. One of our goals is to analyze the thermodynamics of heterotrimer formation involving ANK and MAML binding to CSL. We show that under our ITC conditions, this complex is an obligate heterotrimer–none of the pairwise binary complexes (CSL:ANK, CSL:MAML, ANK:MAML) are populated to appreciable levels. We develop an analytical model that is capable of fitting ITC data where a single component (e.g., MAML) is titrated into a solution of the other two components (e.g., CSL and ANK). This provides, for the first time, a measure of the affinity of the CSL:MAML:ANK ternary complex. In addition to identifying concentrations under which this complex assembles, we can determine whether prebinding RAM to CSL *in trans* affects the free energy of binding for ANK and MAML to CSL, quantitatively testing the potential for allosteric coupling between these two binding sites. Moreover, by comparing the free energy of formation for the CSL:MAML:ANK ternary complex with the free energy of binding for the MAML and ANK domains to CSL *in cis* starting from a prebound RAMANK:CSL complex, we quantify the free energy and enthalpy contribution of the bivalent linkage between RAM and ANK in binding to CSL. These comparisons provide an estimate of the effective concentration afforded by the disordered RAM linker, and reveal the extent to which bivalency promotes cooperative binding of RAM and ANK+MAML to CSL.

## Results

### Experimental design and reaction definitions

In this study, we use ITC to measure the thermodynamics of a series of binding reactions in which parts of the bivalent CSL:RAMANK:MAML ternary complex are formed. Since many of these reactions can be viewed as steps in a larger thermodynamic cycle, comparing the thermodynamics of binding for various reactions allows us to answer questions related to allostery, cooperativity, and bivalency. To describe these complexes compactly and unambiguously, we use a shorthand nomenclature (Figure 1C) in which single letters identify protein segments and domains (C for CSL, M for MAML1, R for Notch1 RAM, and A for the Notch1 ANK domain). For the RAMANK polypeptide, parenthesis indicate that a region or domain (RAM or ANK) is noncovalently tethered to CSL but is not directly engaged with its binding site on CSL.

For example, CR(A) indicates a dimeric complex where RAMANK is directly bound to CSL through its RAM region, but the ANK domain is dissociated (Figure 1C).

With this framework, we can describe reactions that can be probed using ITC (Table 1):

**Table 1:** Reaction definitions for CSL:RAMANK:MAML complexes.

| # | Reaction | $K$ | $\Delta G^\circ$ |
| --- | --- | --- | --- |
| 1 | $C + R \rightleftharpoons CR$ | $K_{CR}$ | $\Delta G_{CR}^\circ$ |
| 2 | $C + RA \rightleftharpoons CR(A)$ | $K_{CR(A)}$ | $\Delta G_{CR(A)}^\circ$ |
| 3 | $C + A \rightleftharpoons CA$ | $K_{CA}$ | $\Delta G_{CA}^\circ$ |
| 4 | $C + M \rightleftharpoons CM$ | $K_{CM}$ | $\Delta G_{CM}^\circ$ |
| 5 | $M + A \rightleftharpoons MA$ | $K_{MA}$ | $\Delta G_{MA}^\circ$ |
| 6 | $C + M + A \rightleftharpoons CMA$ | $K_{CMA}$ | $\Delta G_{CMA}^\circ$ |
| 7 | $CR + M + A \rightleftharpoons RCMA$ | $K_{RCMA}$ | $\Delta G_{RCMA}^\circ$ |
| 8 | $CR(A) + M \rightleftharpoons CMRA$ | $K_{cis}$ | $\Delta G_{cis}^\circ$ |
| 9 | $CR(A) + M + RA \rightleftharpoons R(A)CM(R)A$ | $K_{tetra}$ | $\Delta G_{tetra}^\circ$ |

The first five reactions are bimolecular, whereas the next two (reactions 6-7) are termolecular. Reaction 8, where MAML binds to a complex in which RAMANK is prebound to CSL (CR(A)), is analogous to reaction 6, but the ANK domain binds to CSL *in cis*. Thus, reaction 8 is bimolecular. Reaction 9 starts with the same prebound CR(A) complex as reaction 8, but the ANK binding to CSL is *in trans* rather than *in cis*. Thus reaction 9 is termolecular. The experimental setup used to characterize reactions 1-7 by ITC is shown in Table S1, indicating which protein is in the syringe and which protein (or proteins, for reactions 6 and 7) are in the cell.

For bimolecular reactions 1-5 there is a single polypeptide in the cell and another in the syringe. For termolecular reaction 6 there are two different polypeptides in the cell which do not assemble in isolation. For reaction 7 there are three polypeptides in the cell, but the reaction is termolecular because RAM binds tightly to CSL (and is at stoichiometric excess) prior to injection of MAML.

### RAM binds CSL robustly and independently of ANK

The RAM region of Notch has long been recognized to form a stable binary complex with CSL in solution. This interaction was first identified through yeast two-hybrid and *in vitro* pull-down experiments by Honjo and coworkers [33]. Nam and coworkers mapped the interaction to the central BTD domain of CSL [26]. Kovall and Hendrickson identified a putative binding site for the ΦWΦP motif of RAM on the BTD domain of a crystal structure of CSL from *C. elegans* [22]. ITC studies showed a robust interaction between a peptide spanning the ΦWΦP motif and the BTD domain of CSL [23]; alanine-scanning mutagenesis showed the ΦWΦP motif to be a major determinant of binding energetics, with modest contributions from moderately conserved nearby residues [18]. Subsequent ITC showed that the RAM region binds to larger CSL constructs containing all three domains (NTD, BTD, CTD) with similar, albeit slightly higher affinities than to the isolated BTD domain [12, 31].

The binding of human Notch1 RAM to human CSL by ITC is shown in Figure 2A. As with previous studies, the integrated heats associated with each injection are well-fitted by a heterodimer association model. The model includes an adjustable parameter for the fraction of binding-incompetent material in the injection syringe. The value of 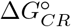 determined from a global fit of the three curves in Figure 2A (−10.2 ± 0.1 kcal/mol) is close to previously reported values (−10.5 kcal/mol from [31], and −10.4 kcal/mol using murine proteins from [12]).

**Figure 2:**
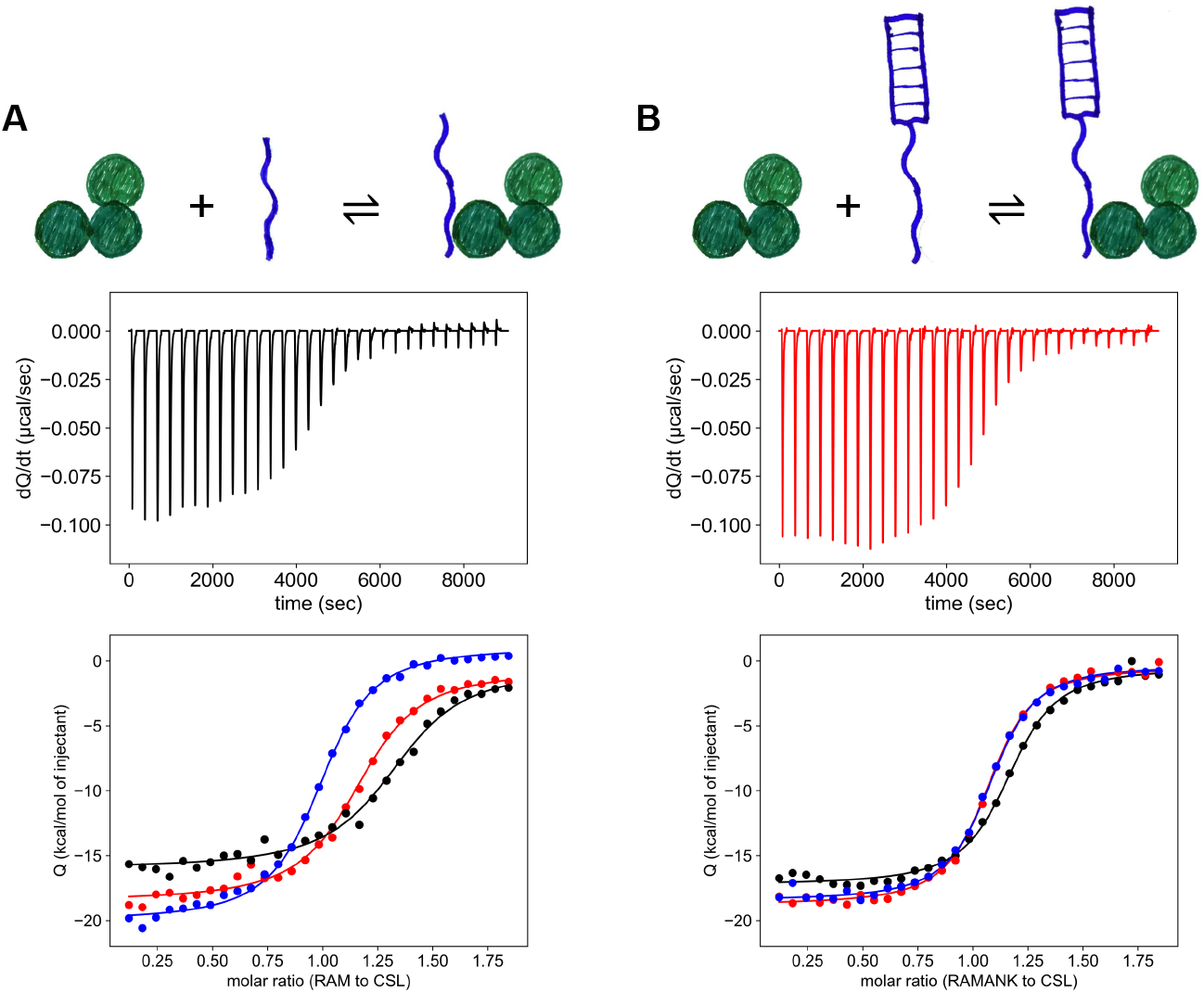
Binding of RAM and RAMANK to CSL measured by ITC. (**A**) RAM, and (**B**) RAMANK titrated into 2 *µM* CSL. Middle panels show baseline-corrected thermograms. Bottom panels show integrated heat values from each injection. The three curves for the RAM titrations (A) are replicate titrations at 20, 23.5, and 27 *µ*M RAM in the syringe (black, red, and blue). The three curves for the RAMANK titrations are replicate titrations each at 20 *µ*M RAMANK in the syringe. The lines show a global fit of the three curves; Δ*G*° and Δ*H*° values are fitted globally, whereas heats of dilution and competent ligand fractions are fitted locally. Competent fractions for RAM (**A**) are 76, 75, and 77 percent (black, red, and blue) and for RAMANK (**B**) are 89, 97, and 96 percent (black, red, and blue).

The binding of RAMANK to CSL is shown in Figure 2B. Again, data are well-fitted by a heterodimer association model. The value of 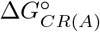 determined from global fit from the three curves in Figure 2B (−10.5 ± 0.1 kcal/mol) is close to a previously reported value (−10.7 kcal/mol from [30]). Moreover, fitted 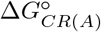 and 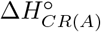 values for RAMANK (Figure 2B) are nearly identical to 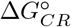 and 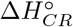 values for RAM (Figure 2A), suggesting that the ANK domain does not interact with CSL when it is tethered to RAM in the absence of MAML. This interpretation is consistent with studies of chimeric human RAMANK orthologues interacting with CSL, which indicate that RAMANK binding to CSL is insensitive to the identity of the ANK domain [28]. Although it is possible that the modest decrease in 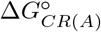 compared to 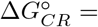 (−10.5 versus −10.2 kcal/mol, Table S1) and a similar enhancement reported in both [4] and [31] reflects a weak CSL-ANK interaction when ANK is tethered *in cis* through RAM, these differences are small, and nearly within error.

### Neither ANK nor MAML bind detectably to CSL

In contrast to the robust interaction of RAM with CSL, neither ANK nor MAML shows evidence of forming a heterodimeric complex with CSL on their own, showing small uniform heat peaks when titrated to a two-fold molar excess relative to CSL (Figure 3A,B). Since these peaks are similar in profile and magnitude to peaks obtained when water is injected into water (Figure 3D), these peaks are likely to be heats of injection. The limit of detection in these experiments can be estimated by a c-value[37], where the c-value is related to the total concentration of CSL in the calorimeter cell, *C*_*t*_, and the dissociation constant by:

**Figure 3:**
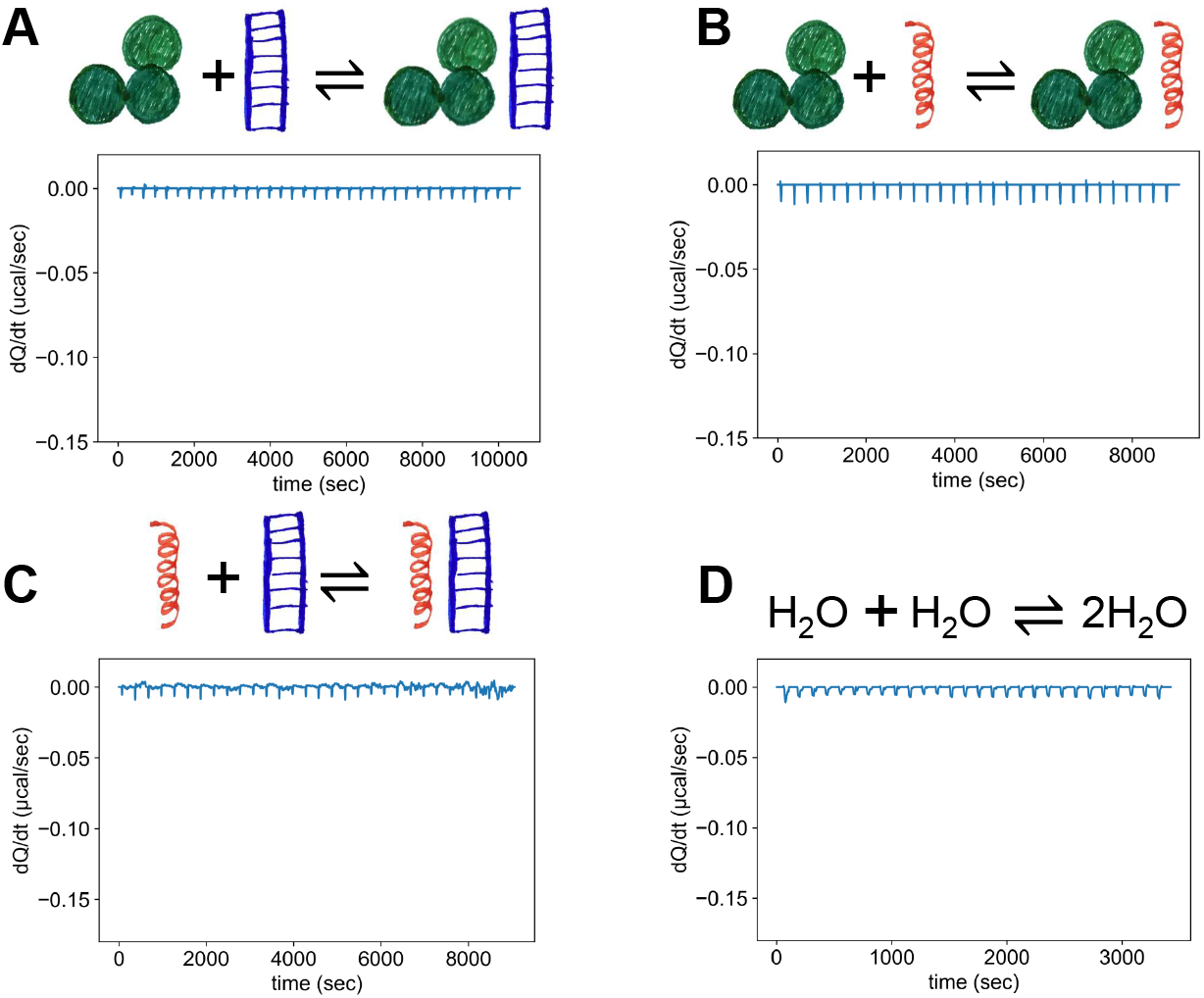
ITC cannot detect formation of binary complexes from ANK, MAML, and CSL. (**A**) 2 *µM* CSL titrated with 20 *µM* ANK. (**B**) 2 *µM* CSL titrated with 20 *µM* MAML. (**C**) 2 *µM* MAML titrated with 20 *µM* ANK. (**D**) Titration of water into water. Plots show baseline corrected thermograms.

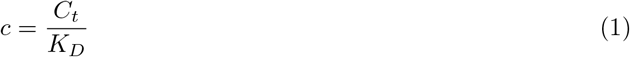

The smallest value of c for which binding can be reliably detected is c = 0.1; since the concentration of CSL in the experiments in Figure 3A,B is *C*_*t*_ = 2 *µM*, the dissociation constant (*K*_*D*_) for each heterodimer must be greater than 20 *µM*.

Similarly, small uniform heat peaks are obtained when ANK is titrated with MAML to a two-fold excess, suggesting that ANK and MAML do not interact in isolation (Figure 3C). These results are consistent with findings from Nam and coworkers [26] that suggest that human Notch1 ANK and MAML1 bind weakly to CSL in isolation. In contrast, Kovall and coworkers were able to detect binding of Drosophila Notch ANK domain to Suppressor of Hairless (the Drosophila CSL orthologue) by ITC, with a 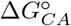 value of −8.4 kcal/mol (a *K*_*D*_ of 0.7 *µM*) [8], suggesting there may be considerable variation in affinity among orthologues.

### CSL + ANK + MAML form a robust complex

We showed in Figure 3A that titration of CSL with ANK gives no heat peaks, but a titration of CSL and MAML with ANK does give clearly discernible heat peaks (Figure S1A). Similarly, Figure 3B showed that CSL titrated with MAML yields no heat peaks; but titrations of CSL and ANK with MAML yield heat peaks (Figure S1B). Titrations of CSL with either ANK or MAML result in no heat peaks, and titrations of MAML with ANK give no heat peaks (3C), but titrations of both MAML and ANK with CSL give heat peaks (Figure S1C). Titrations involving all three proteins (CSL, ANK, and MAML) result in heat peaks regardless of which protein is in the syringe of the calorimeter (Figure S1A-C.) This provides strong evidence for higher-order complex formation.

### A model for studying obligate heterotrimer formation by ITC

Lack of heterodimer formation among CSL, ANK, and MAML (Figure 3) and formation of heat peaks during titrations with all three proteins regardless of which protein is in the calorimeter syringe (Figure S1) suggests that the CSL:MAML:ANK ternary complex is an obligate ternary complex, in which MAML and ANK stabilize one another in their interaction with CSL. To analyze the thermodynamics of CSL:MAML:ANK ternary complex formation, a model is needed that expresses the amount of ternary complex formed in each injection. As with binary complex formation, the heat associated with the *i*^*th*^ injection is equal to the number of moles of CMA (CSL-MAML-ANK) complex formed as a result of injection, *n*_*i*_, times the molar enthalpy of complex formation:

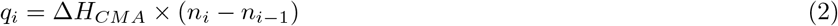

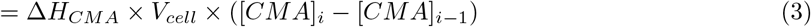

In equation 3, the number of moles of complex is expressed using the product of the molarity ([*CMA*]) and cell volume (*V*_*cell*_).

In principle, the molarity of CMA can be determined from the equilibrium constant, which is logarithmically related to the free energy for formation of CMA 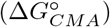. Rearranging the mass-action expression for *K*_*CMA*_ gives

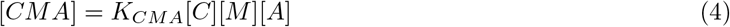

However, none of the free concentrations on the right-hand-side of equation 4 are known (nor is the equilibrium constant, which is to be determined in a fit to the data). A more useful equation can be obtained by replacing the unknown free concentrations [*C*], [*M*], and [*A*] with the total concentrations of each species, which are known^2^. For each species, the free concentration is equal to the total concentration minus the concentration of the ternary complex (e.g., for MAML, [*M*] = [*M*]_*t*_ − [*CMA*]). Replacing each concentration term in equation 4 gives

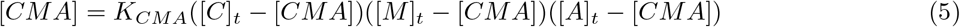

Equation 5 provides a relationship between the concentration of the ternary complex, [*CMA*], and the equilibrium constant, using known total concentrations. However, this equation does not have an explicit form needed for fitting (i.e., [*CMA*] = *f*(*K*_*CMA*_, [*C*]_*t*_, [*M*]_*t*_, [*A*]_*t*_). Rather, equation 5 is cubic in [*CMA*]. Although the cubic equation has an analytical solution, compared to the quadratic equation associated with heterodimer formation, the solution to the cubic equation is quite lengthy and presents some challenges for fitting. A workable solution that generates values of [*CMA*]_*i*_ (the concentration of the ternary complex after the *i*^*th*^ injection) as an explicit function of *K*_*CMA*_ and the starting concentrations ([*C*]_0_, [*M*]_0_, and [*A*]_0_) is presented in Appendix 1. This solution can be inserted into equation 3, which can be fitted to the integrated heats of injection using nonlinear least squares.

### CSL, ANK, and MAML bind robustly as an obligate heterotrimer

In principle, heterotrimer formation among CSL, ANK, and MAML can be measured through six different experimental setups. As demonstrated in Figure S1, any two proteins can be put in the calorimetry cell, and the third protein can be placed in the syringe. Practical considerations favor some setups over others. Because proteins in the syringe need to be at a significantly higher concentration (typically ten-fold molar excess) than proteins in the cell, we chose to put two proteins in the cell and one in the syringe to conserve material (Figure S2A). Since CSL is challenging to prepare at high concentrations compared to ANK and MAML, we put CSL in the cell as the limiting cell ligand. We initially included MAML in the cell at stoichiometric excess to CSL (the excess cell ligand), but during the course of these experiments we learned that the fraction of binding-competent MAML is variable compared to fractions of binding-competent CSL and ANK. Because accurate determination of the incompetent fraction is important to accurately determine binding energetics, and because the incompetent fraction of the excess cell ligand cannot be accurately determined because it has no effect upon the observed molar ratio of syringe ligand to limiting cell ligand (compare Figure S2B,C to S2D), we switched MAML to the syringe. Using this experimental configuration we could accurately fit the energetics of ternary complex formation, without confounding effects from uncertainties in MAML binding competency.

A titration of MAML into CSL and ANK is shown in Figure 4C. At least superficially, the shape of the integrated heat profile resembles that of heterodimer formation. To demonstrate that the heat peaks in this titration result from heterotrimer formation, we collected titrations at the same MAML and CSL concentrations but varying ANK (the excess cell ligand) concentrations (Figure 4E). The titrations at higher ANK concentrations have similar overall heats and midpoints, but steeper transitions. As shown in Figure S2D, this increase in steepness as the excess cell ligand concentration increases is expected for obligate heterotrimer formation. This sensitivity to the concentration of the excess cell ligand, along with the observation that CSL, ANK, and MAML are unable to form heterodimeric species, supports an obligate heterotrimer mechanism for titrations of CSL, ANK, and MAML (Table 1, reaction 6).

**Figure 4:**
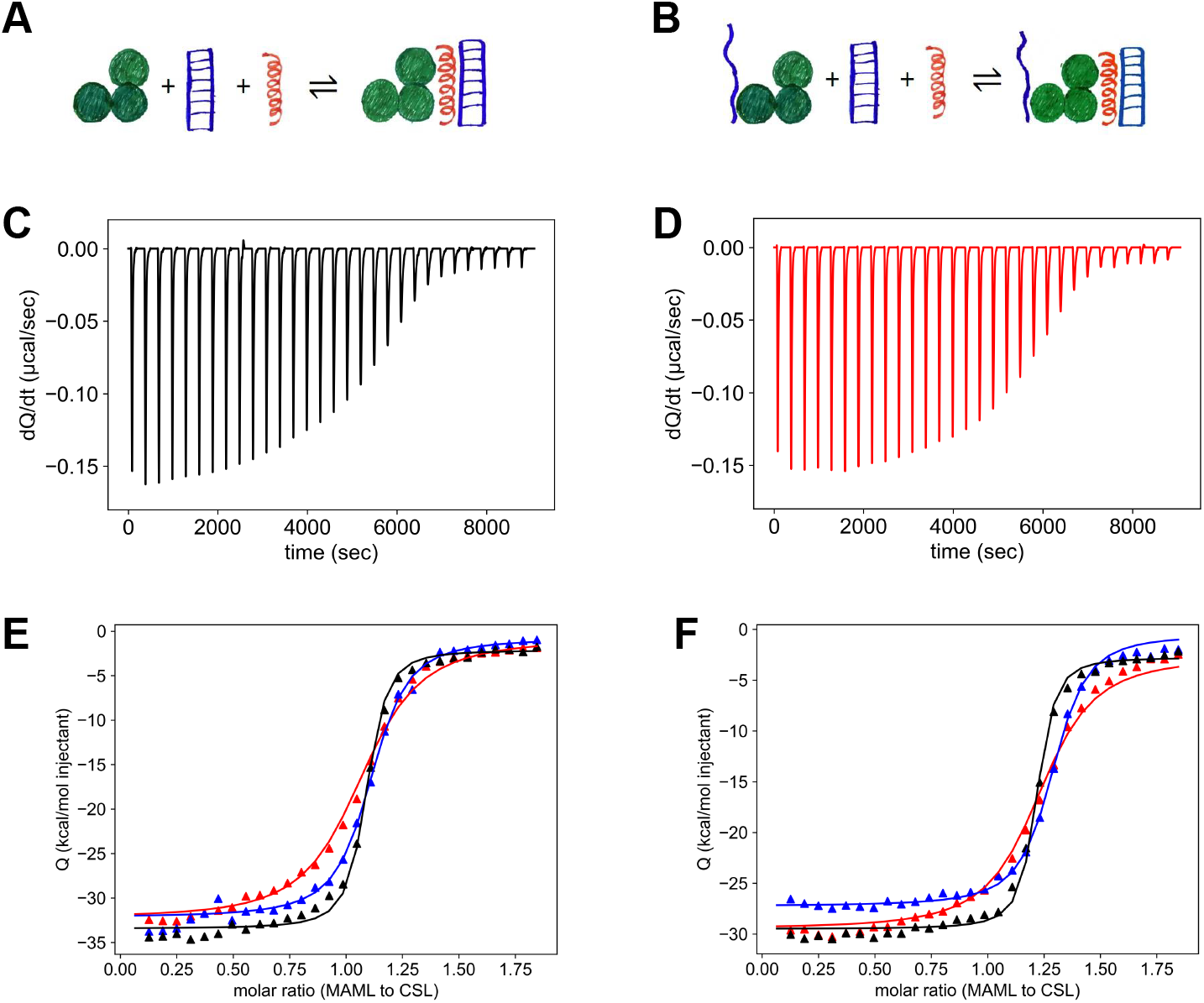
Ternary complex formation involving CSL, ANK, and MAML. Panel (**A**) shows the proteins being titrated for panels (**C**) and (**E**), and panel (**B**) shows the same for panels (**D**) and (**F**). Panels and (**D**) show baseline-corrected thermograms. In both (**C**) and (**D**), 2 *µM* CSL and 8 *µM* ANK in the cell are titrated with 20 *µM* MAML in the syringe. In (**D**) the cell also contains 10 *µM* RAM (*in trans* to ANK), a five-fold stoichiometric excess to CSL. At these concentrations and experimental conditions, CSL should be 99.75 percent saturated with RAM. Panel (**C**) had 78% binding competent MAML, and panel **(D)** had 76% binding competent MAML. Panels (**E**) and (**F**) each show integrated heats of injection for three titrations, wherein both have 20 *µM* MAML in the syringe, 2 *µM* CSL in the cell, and 4 *µM* (red), 8 *µM* (blue), or 22 *µM* (black) ANK respectively. Additionally, Panel (**F**) has 10 *µM* (red, blue, and black) RAM in the cell. MAML competent fractions in (**E**) are 95% (red), 94% (blue), and 96% (black). MAML competent fractions in (**F**) are 82% (red), 80% (blue), and 85% (black). Solid lines are global fits using the obligate heterotrimer model (equation 3).

The heterotrimer model fits well to the individual titrations of MAML into CSL and ANK. To more accurately determine 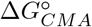 and 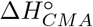, we globally fit the heterotrimer model to three ITC experiments with different ANK concentrations. The global fit accurately reproduces the three titrations, and captures the increase in slope that results from increasing the total concentration of the excess ligand ANK in the calorimeter cell (curves, Figure 4E).

The free energy and enthalpy of formation for the ternary complex between CSL, MAML, and ANK obtained from this global fit 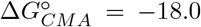 and 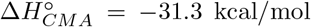, Table S1, reaction 6) are considerably larger in magnitude than typical values for bimolecular protein-protein interactions (e.g.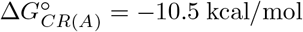 and 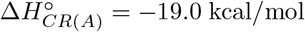). The large negative enthalpy is not surprising, given the considerable surface area that is buried in the CSL:MAML:ANK ternary complex (7,663 Å^2^) [11, 7], and the coil-to-helix transition that is likely to accompany MAML binding [29]. The buried surface area can be used to calculate the heat capacity change expected for CSL-ANK-MAML association [15]. This calculated value (Δ*Cp*_*CMA*_ = −1.58 kcal/mol K) is close to an experimental value (Δ*Cp*_*CMA*_ = −1.46 kcal/mol K) we determined from fitting the temperature dependence of 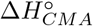 (Figure S3). This agreement is further evidence that the complex formed in titrations of MAML into CSL and ANK is heterotrimeric.

The large negative free energy for ternary complex formation is due, in part, to the stoichiometry of the reaction, which involves three unbound reactants (CSL, MAML, and ANK). 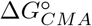 is the free energy of reaction at the standard state, which is defined as one molar concentrations of reactants and products. At this high (and blatantly unphysiological) standard-state, the 3:1 stoichiometry provides strong driving force to form the ternary complex. However as described in the discussion, at concentrations of CSL, MAML, and ANK closer to those used in the ITC experiments, the value of Δ*G*_*CMA*_ increases to a more commonly-encountered, yet still favorable, value.

### Binding of RAM and MAML+ANK to CSL are uncoupled

The binding of RAM to the BTD of CSL has been proposed to allosterically enhance the binding of ANK and MAML to the NTD and CTD of CSL, based on crystallographically-observed structure changes as well as gel-shift assays [12]. The ability to measure the free energy of ANK-MAML interaction with CSL provides a direct way to quantify the thermodynamics of this proposed allosteric coupling, by pre-forming the RAM:CSL complex and measure the free energy of MAML-ANK binding (reaction 7, Table 1) using the ternary fitting model described above. If RAM binding to CSL enhances the ANK-MAML interaction, the standard state free energy of binding ANK and MAML to the pre-formed RAM:CSL complex should be more negative than that for forming the ternary CMA complex in the absence of RAM (reaction 6, Table 1).

Titration of CSL (prebound with RAM) and ANK with MAML (reaction 7, Table S1) produces a heat profile very similar to the analogous titration in the absence of RAM ((reaction 6, Table S1); compare Figures 4C,E and D,F). In these titrations, the cell contains 2 *µM* CSL, 10 *µM* RAM, and varying concentrations of ANK. Based on the free energy of RAM-CSL binding (Table S1, reaction 1), CSL should be 99.8 percent bound with RAM at these concentrations. Globally fitting the data in Figure 4F with the model for obligate ternary complex formation gives a very similar free energy (−18.3 kcal/mol) to that fitted for heterotrimer association without RAM (−18.0 kcal/mol) (Table S1). This result indicates that the binding of MAML and ANK to CSL are not thermodynamically coupled to the binding of RAM in the absence of the covalent linkage between RAM and ANK. These results appear to be at odds with previous results from Friedmann and Kovall using gel-shift assays that show that MAML and ANK bind more readily to CSL in the presence of RAM [12]. One difference between these two studies is that the present study uses human proteins whereas the Friedmann study used *C. elegans* and mouse proteins. Another difference is that in the Friedmann study, CSL was bound to DNA. It is possible that coupling between the RAM- and MAML-ANK binding sites on CSL is mediated by bound DNA; however the DNA binding specificity of CSL was previously found to be unaffected by binding CSL with RAMANK or RAMANK+MAML [9]. Consistent with a DNA-agnostic interpretation, the Friedmann study found the affinity between CSL and RAM to be within error of the affinity between CSL+DNA and RAM.

### Binding of ANK and MAML to CSL *in cis* and the energetics of bivalency

The ability to separate the CSL:RAM and CSL:ANK:MAML interactions (Table S1 reactions 1, 2, and 6) and quantify their binding energies provides a means to extract the contribution of bivalency from the overall assembly energy of the bivalent ternary complex (CMRA in Figure 1C). Since the binding of RAM and ANK/MAML to CSL *in trans* seem to be uncoupled (Figure 4), the binding energy of the bivalent ternary complex is equal to the sum of the separate CR(A) and CMA binding energies plus the free energy of bivalent coupling between the two 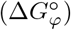, i.e.,

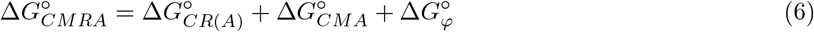

A simple way to measure the free energy of bivalent coupling is to compare the assembly energy of the ternary CSL:MAML:ANK complex (reaction 6, Table S1) with the analogous reaction starting from a prebound CSL:RAMANK complex (reaction 8, Table S2). The binding energy of MAML to the prebound CR(A) complex should include the contribution from binding of ANK, since ANK does not appear to be bound in the CR(A) complex (Figure 2). The free energy of MAML binding to CR(A) *in cis* is equal to the assembly energy of the CMA complex plus the free energy of bivalent coupling, 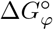:

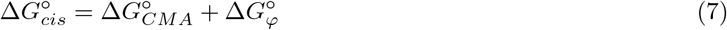

Rearranging this equation leads to an experimentally accessible definition of the free energy of bivalent coupling:

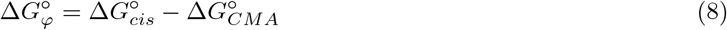

Similarly, the enthalpy of bivalent coupling is

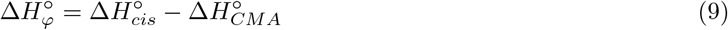

Five titrations of CSL prebound by RAMANK (CR(A)) with MAML are shown in Figure 5. These titrations differ in the amount of RAMANK in the cell along with 2 *µM* CSL, though in all cases, RAMANK is in excess of CSL and should be sufficient to fully saturate the CR(A) complex. Unlike the titration of CSL + ANK with MAML (reaction 6, Figure 4), the five heat profiles all have the same slope, consistent with the expected heterodimer mechanism.

**Figure 5:**
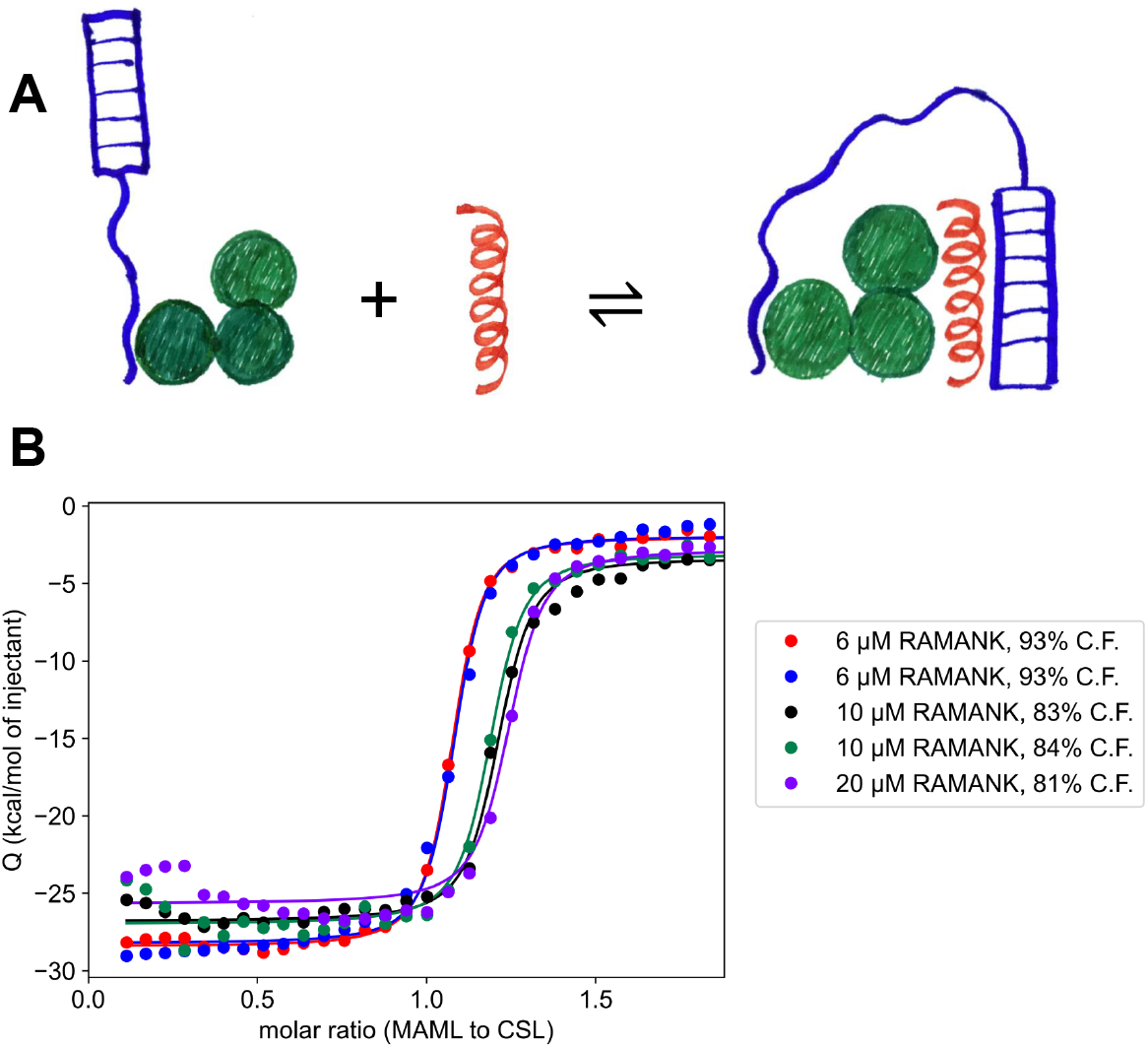
Binding of MAML and the ANK domain *in cis* to CSL, using a heterodimer model to determine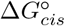. Panel (**A**) shows the reaction being considered, in which MAML is titrated into CSL prebound with RAMANK. Panel (**B**) shows integrated heats of injection from five titrations with RAMANK at concentrations varying from 6 *µM* to 20 *µM* prebound to 2 *µM* CSL in the ITC cell, titrated with 20 *µM* MAML (locally fitted binding competent fractions of MAML are abbreviated as C.F. above). At these concentrations, CSL should be > 99% percent saturated with RAMANK. Curves are from globally fitting the data for 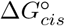 and 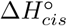 using the heterodimer model shown in (**A**).

Globally fitting the data in Figure 5 to a bimolecular association scheme leads to an estimate of 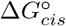 of − 11.6 kcal/mol and 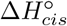 of −28.3 kcal/mol. Combining these values with the free energy and enthalpy of forming the ternary complex (reaction 6, Table S1) with equations 8 and 9 gives the free energy and enthalpy of bivalent coupling (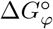 and 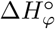) of +6.4 kcal/mol and +3.0 kcal/mol. Although the positive value of 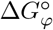 might seem to suggest an energetically unfavorable reaction, this interpretation is complicated by the stoichiometry difference between reactions 6 and 8 and the use of a one molar standard state. As developed below, at more dilute (physiologically relevant) concentrations, this value of 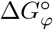 produces considerable coupling between RAM and ANK/MAML binding to CSL.

### A more complete model to determine the energetics of bivalency

In addition to forming the bivalent ternary (CMRA) complex, the MAML titrations of CSL prebound with RAMANK (Figure 5) have the potential to form a tetrameric complex containing two RAMANK molecules (R(A)CM(R)A, Figure 1C). This tetrameric complex, which would be favored at high RAMANK concentrations^3^, is not accounted for in the model used to fit 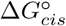 in the previous section. Neglecting tetramer formation could adversely affect fitted parameters, in particular, the parameters associated with bivalency (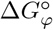 and 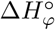).

Here we develop a fitting model that includes all species that can form in a mixture of CSL, RAMANK, and MAML, including CR(A), CMA, and R(A)CM(R)A, in addition to the bivalent ternary complex (CMRA). Derivation of an analytical fitting equation that includes the tetrameric species is significantly complicated by the quartic equation that results from expression of *K*_*tetra*_ in terms of total concentrations. Instead, we have developed a routine that uses numerical methods to solve for the unbound concentrations needed to fit the ITC data with nonlinear least squares (Appendix 3). This approach is based on that of Vega et al. [35]. To simplify analysis, we fixed Δ*G*° and Δ*H*° for CR(A) and CMA at values determined in Figures 2 and 4 (Table S1), and fit 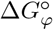 and 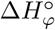 in titrations of preformed CR(A) complexes with MAML (Figure 6).

**Figure 6:**
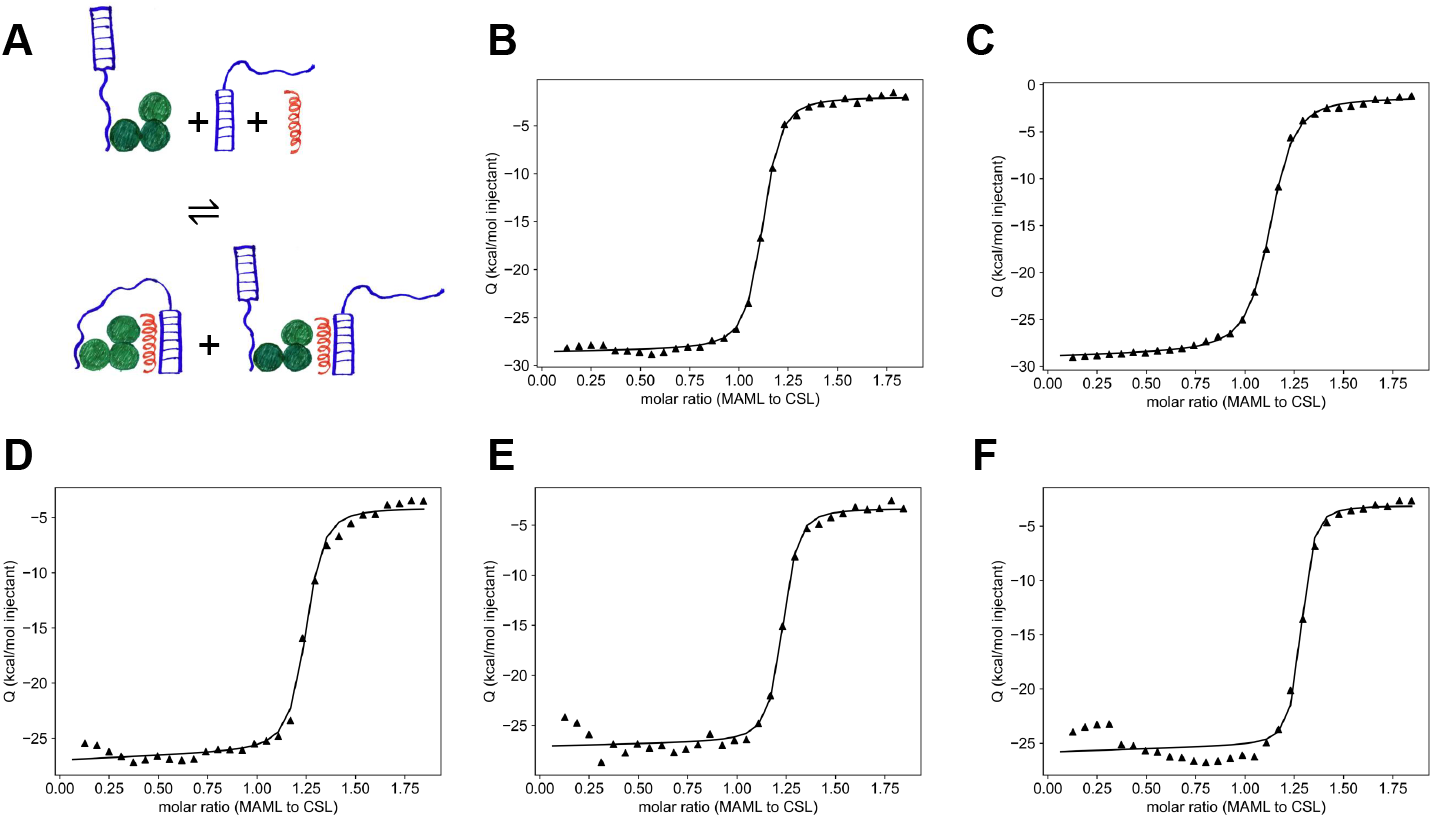
Numerical fitting to determine 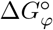 from titrations of prebound CSL:RAMANK with MAML. (**A**) Highest-populated ligation states during the experiment. In all five titrations, MAML is in the syringe at 20 *µM*, and CSL is in the cell at 2 *µM* with a molar excess of RAMANK at 6 *µM* (B and C), 10 *µM* (D and E), and 20 *µM* (**F**). At these concentrations, CSL should be > 99.8 percent saturated with RAMANK. Fitted binding competent fractions of MAML are 93%, 93%, 83%, 84%, and 80% (B-F). Curves are the result of numerical fits for 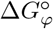 and 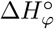 (Appendix 3).

Five titrations of CSL prebound with different concentrations of saturating RAMANK with MAML are shown in Figure 6. Individually fitting each curve using the numerical solution described in Appendix 3 gives a 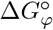 value of +6.5 ± 0.2 kcal/mol and a 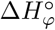 value of +9.5 ± 1.5 kcal/mol (Table S2). This value of 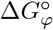 is within error of the value determined using equations 8 and 9 (Figure 5). However, there is a significant change in 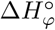, which increases from 3.0 to 9.5 kcal/mol when tetramer is accounted for. This change in the value of 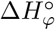 leads to a favorable entropy of bivalency (+10.1 cal/mol K).

## Discussion

The assembly of the Notch transcriptional activation complex is a critical step in the Notch signaling pathway, and is essential for development and homeostasis. The bivalent interaction between the RAM and ANK regions of the Notch intracellular domain and CSL has been appreciated for more than two decades, first from functional and pull-down studies and then from structural studies [16, 26, 4]. Transcriptional activation studies have shown that maximal activity requires RAM and ANK on the same polypeptide, as opposed to being on two separate polypeptides (which are incapable of the bivalent interaction) [17]. The mechanisms underlying bivalency have been discussed extensively, and include allosteric coupling between the RAM and ANK binding sites [12], and an effective concentration enhancement by which binding of RAM (or ANK via MAML) increases the local concentration of ANK (or RAM) near its site on CSL [3]. There has been some support for enhancement of effective concentration from deletion and insertion studies studies within the linker [32], but experimental determination of the thermodynamic contributions of bivalency and the magnitude of the effective concentration enhancement have been lacking. By developing an ITC assay to measure ternary complex formation along with an equation for data analysis, we have been able to provide the experimental thermodynamic quantities needed to determine the effective concentration enhancement.

### The free energy of CSL:MAML:ANK ternary complex formation and the one molar standard state

Although none of the pairwise interactions among CSL, MAML, and ANK can be detected at ITC concentrations (2 − 10 *µM*), a robust and saturable complex is formed at an approximate 1:1:1 stoichiometry when all three proteins are combined, regardless of which protein is in the syringe and which proteins are in the calorimeter cell (Figure S1). Injection heats are well-fitted by a model for ternary complex formation, and show the expected sensitivity to the concentration of the excess cell ligand (Figure 4).

As mentioned above, the free energy for formation of the CSL:MAML:ANK ternary complex 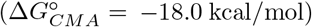 is larger in magnitude than values typically measured in the ITC for bimolecular association. Despite this large value, the titration is shallow enough in the transition region to detect and quantify measurable equilibrium between the ternary complex and the dissociated proteins, as is seen for heterodimer reactions with smaller free energies of binding (see Figure 2 for example).^4^ This difference in free energy is largely a result of the difference between the stoichiometry of the reaction, and the fact that free energies are reported at a one molar standard state.

Although a concentration of one molar reactants and products would strongly favor the termolecular association of CMA due to its 3-to-1 stoichiometry, the free energy falls dramatically as concentration decreases. For example, at concentration of one micromolar reactants and products (i.e., [*C*] = [*M*] = [*A*] = [*CMA*] = 1 *µM*), the free energy of reaction can be calculated from the familiar equation

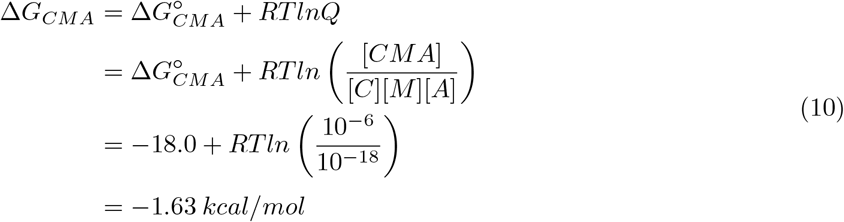

At this concentration, the ternary complex would be favored, but only modestly so. This quantity can be thought of as a free energy of ternary complex formation at a new standard state of one micromolar reactants and products. Although this standard state is unconventional, it is a certainly closer to cellular concentrations than the typical one molar standard state.

The concentration at which CSL is half-saturated with MAML and ANK at equilibrium can be calculated using equation 10. At this concentration, Δ*G* = 0. From equation 10,

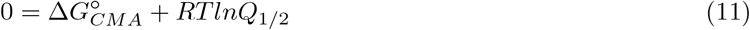

where *Q*_1*/*2_ gives the equilibrium concentration where [*CMA*] = [*C*]. Rearranging gives

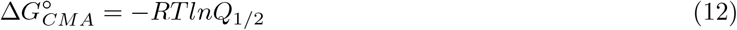

Assuming equal total concentrations (i.e., [*C*]_*t*_ = [*M*]_*t*_ = [*A*]_*t*_), *Q*_1*/*2_ can be expressed as

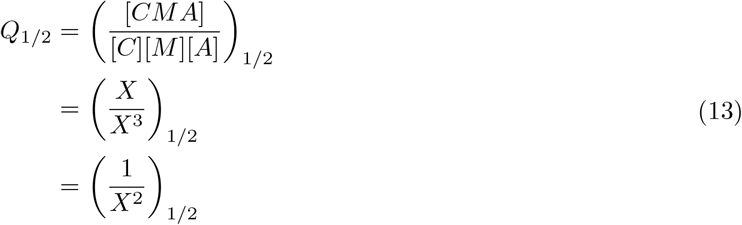

where *X* is the molar concentration that brings about half-saturation. Combining equations 13 and 12 gives

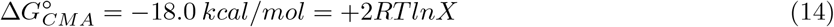

Rearranging gives an free equilibrium concentration for half-saturation (*X*) of 253 *nM*, corresponding to total concentrations of [*C*]_*t*_ = [*M*]_*t*_ = [*A*]_*t*_ = 506 *nM*. Thus, at half-micromolar total protein concentrations, the CMA ternary complex is half-formed. Below this concentration, the CSL:MAML:ANK ternary complex should be predominantly dissociated. Though the concentrations of CSL and MAML in the nucleus are not well known, recent work using Notch2 suggests nuclear concentrations of 2 *nM*[34]. Therefore it seems unlikely that the concentrations of NICD, CSL, and MAML would exceed 506 *nM*, and in the absence of bivalency, the CSL:MAML:ANK ternary complex would not be expected to assemble to high levels in a cellular context.

### Analysis of bivalency in terms of effective concentration

Here we will analyze bivalency by comparing equilibrium constants. Because the dimensions of equilibrium constants are powers of molarity, this analysis provides a concentration-based view of bivalency, and justifies interpreting *K*_*φ*_ (the equilibrium constant associated with 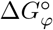) as an effective concentration. To connect *K*_*φ*_ with effective concentration, it is helpful to relate *K*_*φ*_ to the two reactions that define it, namely *K*_*cis*_ (Table 1, reaction 8) and *K*_*CMA*_ (Table 1, reaction 6). Writing the standard state free energies from equation 7 in terms of equilibrium constants,

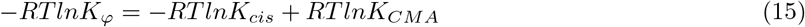

Rearranging gives *K*_*φ*_ as a ratio of binding constants for binding of ANK and MAML to CSL *in cis* and *in trans*:

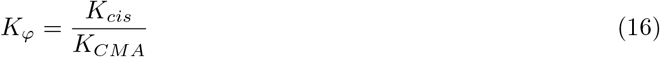

Because *K*_*cis*_ and *K*_*CMA*_ have units of *M*^−1^ and *M*^−2^, *K*_*φ*_ has units of *M* (that is, *K*_*φ*_ is a concentration). Using the value of 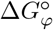 fit in Figure 6, the value of *K*_*φ*_ is 17 *µM*. This value is lower than that reported for the bivalent interaction between retinoblastoma protein and adenovirus early region 1A protein (520 *µM*) [14], an observation that is closely related to the rather large positive value of 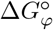.

To demonstrate the relationship between *K*_*φ*_ and effective concentration, we can compare equations that give the fraction of CSL bound with ANK (and its obligate binding partner MAML) *in cis* and with ANK *in trans* (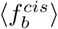 and 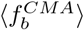). The fraction of CSL bound *in cis* with ANK is

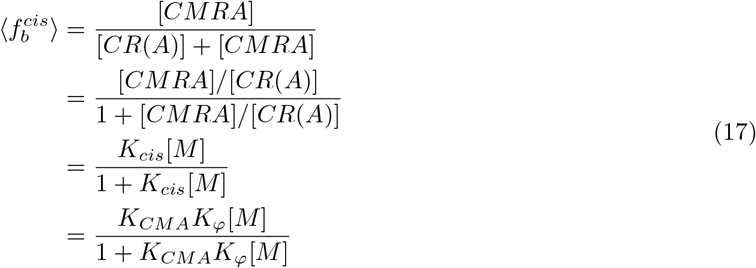

As would be expected for a bimolecular reaction, 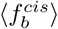 is a simple rectangular hyperbola that depends on ligand concentration (here, MAML).

The fraction of CSL bound with ANK *in trans* is

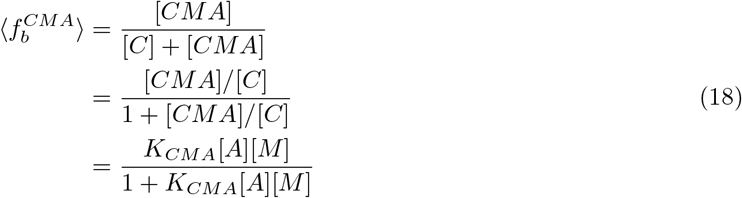

Although 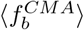 becomes sigmoidal when both [*M*] and [*A*] are increased, it is a rectangular hyperbola in either [*M*] or [*A*] when the other is held constant^5^. This is shown in Figure 7, where ANK is titrated at constant MAML concentrations (solid curves). The ANK concentration that brings about half-saturation decreases with increasing MAML concentration, reflecting an apparent increase in ANK affinity. Indeed, at a fixed MAML concentration, the product *K*_*CMA*_[*MAML*] in equation 18 can be viewed as an apparent equilibrium constant that increases with MAML concentration. This apparent increase in binding affinity with increasing MAML concentration is the basis for the increased steepness of the integrated heat peaks in Figure 4^6^.

**Figure 7:**
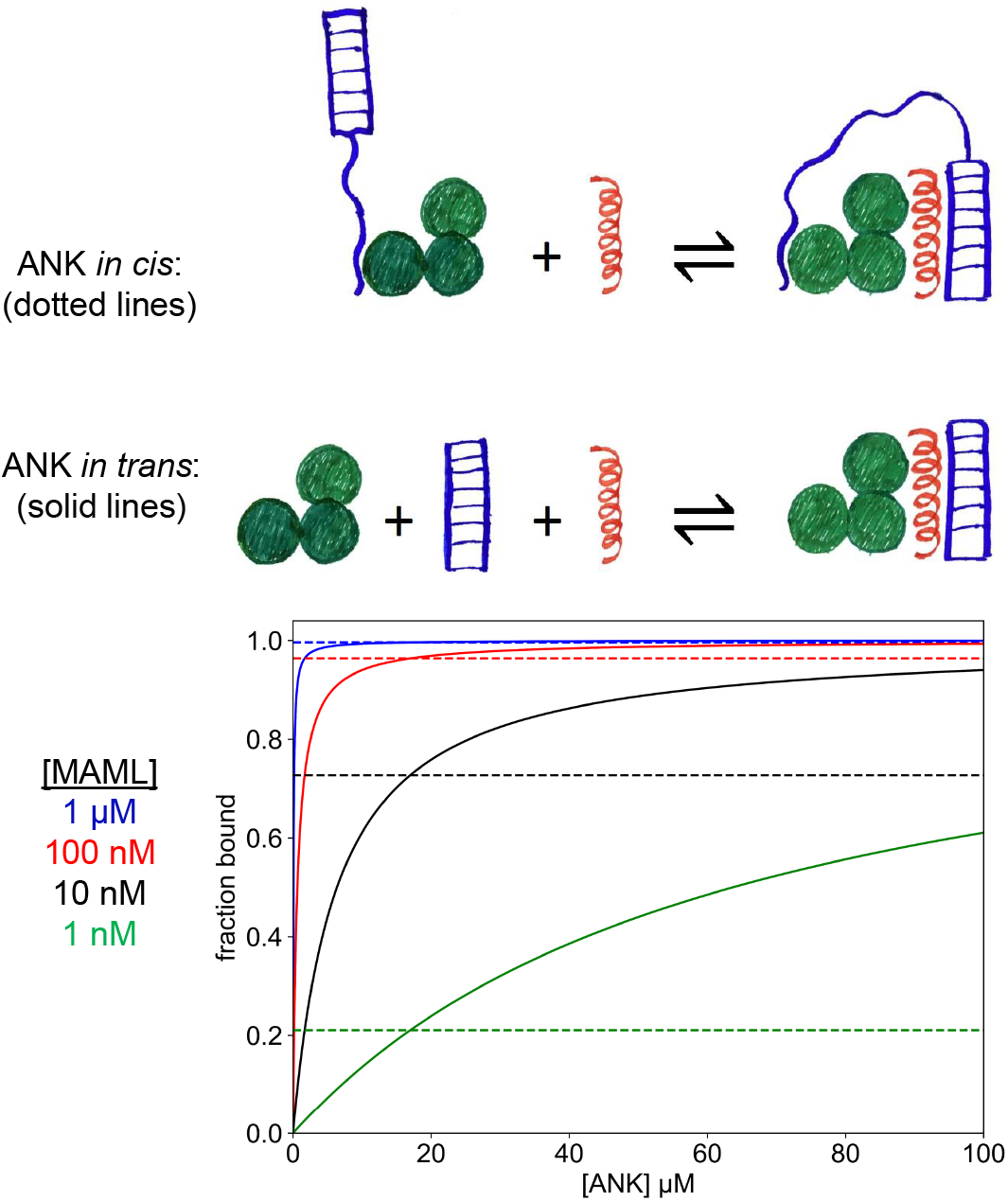
The fraction of ANK bound to CSL *in cis* and *in trans* at constant [MAML] concentrations. When ANK binds *in trans* at constant [MAML] (solid hyperbolas), it does so with an apparent binding constant of *K*_*CMA*_[*MAML*]. When ANK binds *in cis* (dashed flat lines), a constant level of saturation is obtained, because the concentration of ANK is constant. The concentration of ANK where 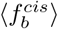 and 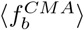 intersect is the effective concentration of ANK when noncovalently tethered to CSL (17 *µM*). Since 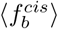 and 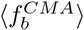 both increase with [MAML], the effective concentration (the intersection point) remains constant.

In contrast, CSL saturation by ANK *in cis* does not depend on ANK concentration (equation 17). At a given MAML concentration, 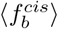 is constant (see the dashed lines in Figure 7), although it increases with MAML concentration in the same way as does 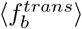. The difference in the ANK dependence of 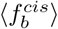 and 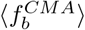 results in a single intersection point between the two curves. Although the saturation value at this intersection point depends on MAML concentration, the ANK concentration at which these two curves intersect (which we will refer to as [*A*]^∗^) does not.

The significance of the intersection point between 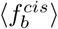 and 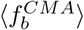 can be appreciated by comparing the expressions for these two quantities at the intersection point. At 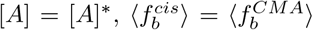. Thus, from equations 17 and 18,

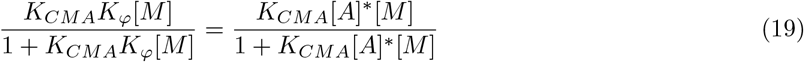

The only difference between the left- and right-hand sides of equation 19 (assuming MAML is at the same concentration on the left and right sides) is that *K*_*φ*_ on the left is replaced with [*A*]^∗^ on the right. Thus, to maintain the equality,

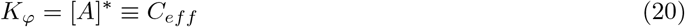

In words, *K*_*φ*_ is the concentration of ANK that would be required to bring about the same saturation with ANK binding CSL *in trans* as is obtained with ANK binding CSL *in cis*. Alternatively, *K*_*φ*_ is the concentration of ANK when tethered by RAM bound to CSL, which is the definition of *C*_*eff*_, the effective concentration. Thus, *K*_*φ*_ = *C*_*eff*_.

### The free energy of bivalent coupling at a dilute standard state

As alluded to in the results, the positive value of 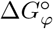 is a result of the difference in stoichiometry of reactions 6 and 8 (from Table 1), and the use of a one molar standard state. Here we will explore how 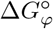 changes when a more dilute (and more physiological) standard state is chosen. In general, a new standard state reaction free energy can be obtained using the equation

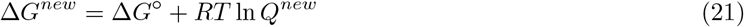

where Δ*G*^*new*^ is the reaction free energy at a new standard state, Δ*G*° is the reaction free energy at the 1 M standard state, and *Q*^*new*^ is the reaction quotient in which all concentrations are simultaneously changed to the new standard state concentration.

For simplicity we will choose a new standard state such that the concentrations of all reactants and products are the same. For the trans reaction,

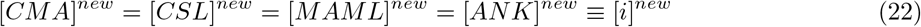

For the bivalent coupling,

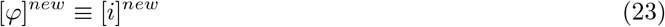

These new standard state concentrations can be inserted into the *Q*^*new*^ term of equation 21 to generate free energies for the trans reaction and for bivalent coupling:

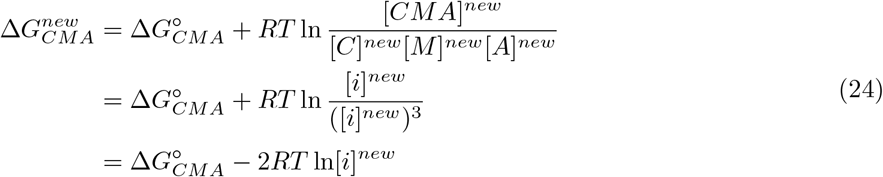

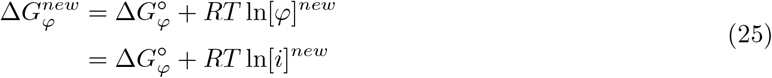

Following from equations 25 and 24, the free energy of the cis reaction at a new standard state can be calculated using equation 7,

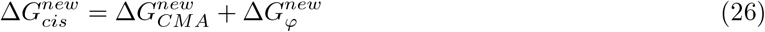

The standard state free energies in equation 26 are plotted as a function of standard state concentration (all reactants and products) in Figure S4. Because 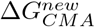 and 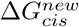 describe association reactions, they increase linearly with decreasing standard state concentration. However, they increase with different slopes due to their different stoichiometries, intersecting at the effective concentration (17 *µM*). Because 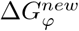 is the difference between 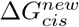 and 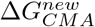 (equation 26), their difference in slope results in a decrease in 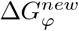 with decreasing standard state concentration, falling to zero at *C*_*eff*_. Below 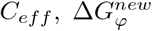 is negative, indicating that the cis reaction (reaction 8, Table 1) is more favorable than the trans reaction (reaction 6, Table 1).

### Bivalency as a mechanism to couple RAM and ANK binding to CSL

Using the experimentally determined values of 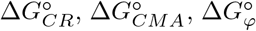 (Tables S1 and S2), we can determine the populations of partly and fully ligated species as a function of protein concentrations using binding polynomials for CSL. This approach provides a means to evaluate how the linkage of RAM and ANK binding to CSL *in cis* promotes bivalent ternary complex formation (CMRA), where both RAM and ANK are bound, over singly bound (CR(A), (R)CMA) ligation states.

Figure 8 shows the populations of different ligation states of CSL at different MAML concentrations when RAMANK is titrated. At high MAML concentration (Figure 8 A), CSL converts from completely unbound to the bivalent CMRA complex where both RAM and ANK are bound to their sites on CSL. The midpoint for this transition is 74 *pM* RAMANK. Populations of singly-ligated species (CR(A) and (R)CMA) are suppressed at all RAMANK concentrations. At high RAMANK concentrations, the bivalent CMRA complex shifts to the 2:1:1 RAMANK:CSL:MAML complex (R(A)CM(R)A, red curves, Figure 8). The RAMANK concentration at which the bivalent CMRA and tetrameric R(A)CM(R)A complexes have equal populations (the point at which the purple and red curves cross, Figure 8) is equal to the effective concentration *C*_*eff*_ (17 *µM*) regardless of the MAML concentration.

**Figure 8:**
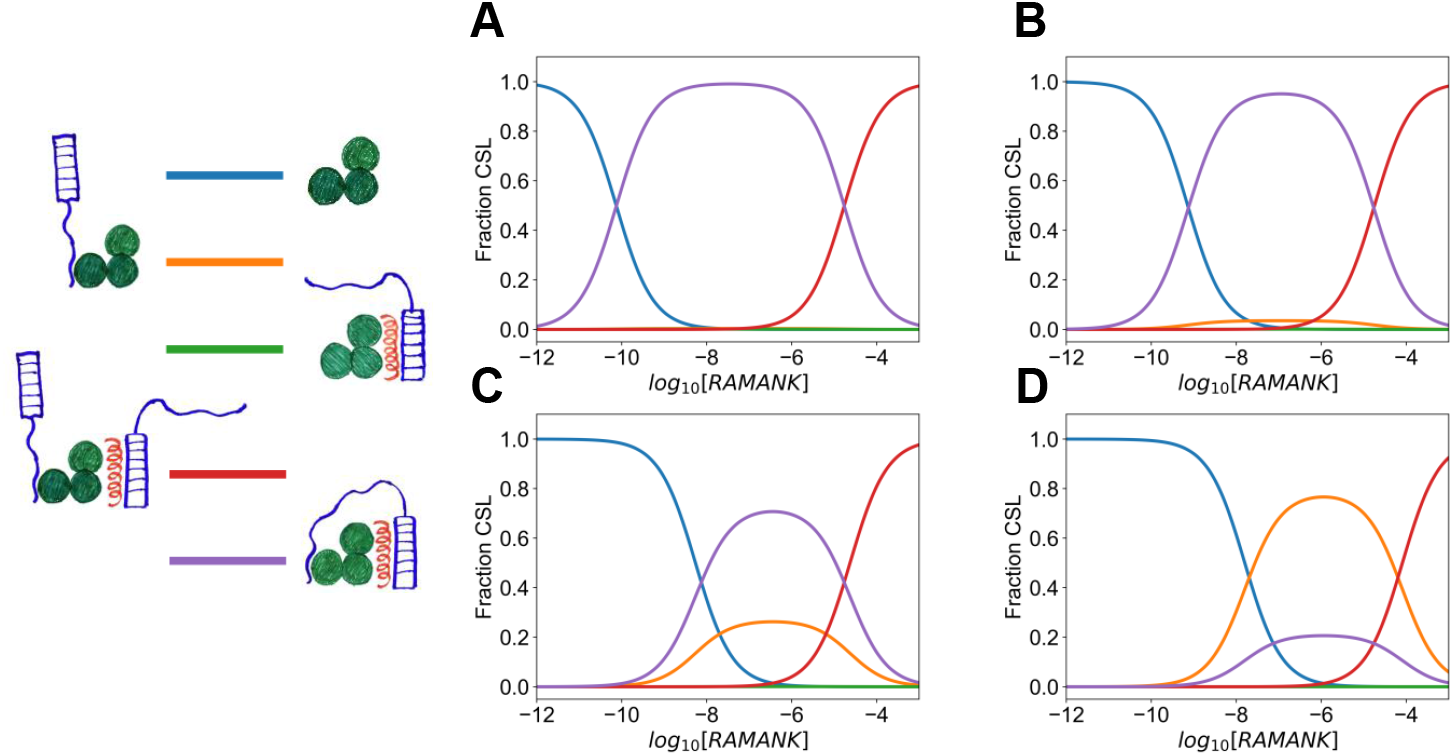
Populations of ligation states of CSL as a function of free (unbound) RAMANK and MAML concentration. The four panels show RAMANK titrations of CSL, MAML concentrations of (**A**) 1 *µM*, (**B**) 100 *nM*, (**C**) 10 *nM*, and (**D**) 1 *nM*. Curves correspond to different ligation states of CSL (see legend), calculated using the binding polynomial (equation 49, Appendix 2) and values of 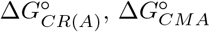, and 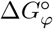 from Tables S1 and S2.

As the concentration of MAML is decreased, the midpoint for conversion from C to CMRA shifts to higher RAMANK concentrations (Figure 8 B,C,D). The magnitude of this shift is inversely proportional to the change in MAML concentration: decreasing MAML by one order of magnitude (1 *µM* to 100 *nM*) increases the midpoint from 74 *pM* to 740 *pM*. Another order of magnitude decrease ([*MAML*] = 10 *nM*) increases the midpoint to 7.4 *nM*, and a further decrease ([*MAML*] = 1 *nM*) increases the midpoint to 74 *nM*. Although the monovalently-ligated species are suppressed at high MAML concentrations, as MAML concentration decreases to the 1-10 *nM* range, the species of CSL bound with RAM but with ANK dissociated (CR(A), orange curve, Figure 8) is formed at appreciable concentrations.

In addition to comparing the different ligation states of CSL in complex with RAMANK, the binding polynomial (Appendix 2) can be used to determine the extent to which bivalency increases RAM and ANK binding, by comparing the occupancy of RAM and ANK at their CSL binding sites when on a single polypeptide (capable of binding to CSL *in cis*) to occupancies when RAM and ANK are on two separate polypeptides (only capable of binding to CSL *in trans*) (Figure 9). When RAM and ANK are on the same polypeptide, occupancy at their respective sites is half-saturated at 74 pM (at 1 *µM* MAML), whereas titration with separate RAM and ANK polypeptides does not half-saturate until 20 and 65 *nM* (Figure 9 A,B). Thus, bivalency increases occupancy at both sites by about three orders of magnitude.

**Figure 9:**
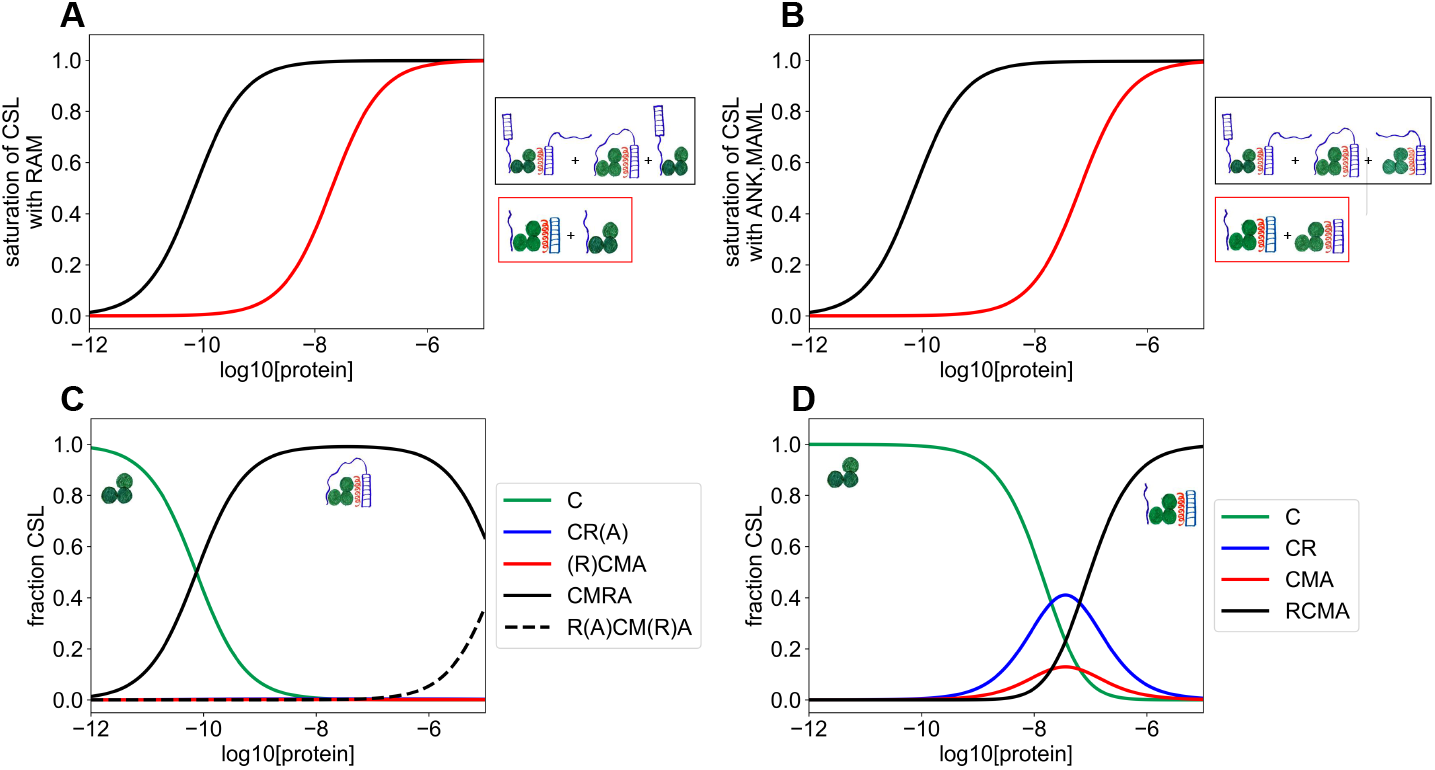
Comparison of CSL saturation with RAMANK (one polypeptide) to saturation with RAM and ANK (two separate polypeptides) at equal concentrations. (**A**) Saturation of CSL with RAM when RAM and ANK are on the same polypeptide (black), and when RAM and ANK are at the same concentrations but on separate polypeptides (red). (**B**) Saturation of CSL with ANK and MAML, when RAM and ANK are on the same polypeptide (black), and when RAM and ANK are on separate polypeptides at the same concentrations (red). Saturation in (**A**) and (**B**) includes all species in which RAM (**A**) and ANK + MAML (**B**) are directly bound to their sites on CSL. (**C**) shows ligation states of CSL with RAMANK (one polypeptide), and (**D**) shows ligation states of CSL with RAM and ANK (two polypeptides). In all four plots, MAML is at a concentration of 1 *µM* and all concentrations are free (rather than total).

Moreover, whereas the ternary bivalent CMRA species is the dominant ligation state when RAM and ANK are on the same polypeptide (Figures 8, 9C), half-ligated states populate significantly when RAM and ANK are two separate polypeptides (Figure 9D). The doubly ligated RCMA state (black curve, Figure 9D) is not half-saturated until 96 *nM* RAM and ANK. This analysis shows that the covalent linkage between RAM and ANK enhances CSL saturation by three orders of magnitude, combining two moderate affinity interactions to facilitate saturation at sub-nanomolar concentrations.

## Methods

### Expression and purification of CSL, RAM, RAMANK, and ANK

Protein samples of CSL, RAM, RAMANK, and ANK were expressed and purified as in [31] and [23], using modified pMAL-c2x expression vectors with N-terminal His_12_ tags and MBP solubility tags. Briefly, expression vectors were transformed into *E. coli* strain BL21(DE3) and grown with Ampicillin at 37°C while shaking at 200rpm to an O.D.600 between 0.6-0.8. Expression was induced by adding IPTG to 1mM and shaking at 200rpm and 20°C for 12-18 hours. Cells were pelleted by centrifugation at 4,424×g and stored at −80°C. For purification, cell pellets were thawed to 20°C and resuspended in 25mM Tris, 50mM NaCl, 0.5mM tris(2-carboxyethyl)phosphine (TCEP), pH8 (lysis buffer) and one c0mplete protease inhibitor tablet (Roche catalog no. 11836170001) per 100 mL lysate. Resuspended cells were sonicated on ice and were centrifuged at 30,966×g for 40 minutes at 4°C. The cleared supernatant was brought to 2m*M MgCl*_2_ and treated with DNaseI and benzonase at 20°C for 1 hour. The sample was brought to 500mM NaCl (now in equilibration buffer; 25mM Tris, 500mM NaCl, 0.5mM TCEP, pH8) and loaded onto a Ni-NTA gravity column at 4°C. The column was washed with equilibration buffer containing 100 mM imidazole, and protein was eluted using equilibration buffer containing 750 mM imidazole. Eluted protein was dialyzed into 25 mM Na Phos, 150 mM NaCl, 0.5 mM TCEP pH7 (ion exchange buffer A) overnight at 4°C in the presence of 2 mg TEV protease. Cleaved sample was passed over a Ni-NTA gravity column at 4°C to remove any uncleaved protein. The Ni-NTA column was washed with ion exchange buffer A containing 10mM imidazole. The flow-through and wash were collected, filtered, and purified using either a HiTrapQ HF anion exchange column or a HiTrapSP HF cation exchange column (Cytiva) by eluting with a linear gradient of 1 mM to 1 M NaCl in 25 mM Na Phos, 0.5 mM TCEP, pH7. Fractions containing pure protein, as judged by SDS PAGE, were combined and dialyzed at 4°C in 25 mM Na Phos, 150 mM NaCl, 0.5 mM TCEP pH7.

### Expression and purification of MAML

Human MAML1 (residues 8-76) with a C-terminal tryptophan for concentration measurement was expressed and purified as in [28] using a pET expression vector with an N-terminal His_6_ tag and a SUMO solubility tag. Briefly, the expression vector was transformed into *E. coli* strain BL21(DE3) and grown with Kanamycin at 37°C while shaking at 200rpm to an O.D.600 of 0.8. Expression was induced as described above for CSL. Cells were pelleted by centrifugation at 5,000×g and resuspended, lysed, and pelleted as described above for CSL. His-SUMO-MAML1-W localizes to the lysate pellet, which was washed twice in 100 mM Tris pH 7.4, 5 mM EDTA, 1 M urea, 0.5 mM TCEP, 1% Triton X-100 followed by a wash in 100 mM Tris pH 7.4 and 0.5 mM TCEP. Pellets were re-suspended using a Dounce homogenizer and re-pelleted by centrifugation at 15,000×g for 25 minutes. The washed pellet was resuspended in 8 M urea, 25 mM Tris pH 8.0, 250 mM NaCl, 0.5 mM TCEP (binding buffer) and batch bound to Ni-NTA resin overnight with rotation at 4°C. The resin was loaded onto a gravity column and washed with binding buffer to remove unbound protein. Protein was refolded while bound to the resin by washing with 25 mM Tris pH 8.0, 250 mM NaCl, 0.5 mM TCEP (refold buffer). His-SUMO-MAML1W was eluted from the resin using refold buffer containing 300 mM imidazole, and the sample was brought to 100 mM EDTA. SUMO protease Ulp1 was added to the His-SUMO-MAML1W, and sample was dialyzed 8 hours at 4°C in 50 mM Tris pH 8.0, 150 mM NaCl, 0.5 mM TCEP, 5% glycerol followed by dialysis overnight at 4°C in 25 mM Na Phos pH 8.0, 50 mM NaCl, 0.5 mM TCEP. Cleaved sample was centrifuged at 15,000×g for 30 minutes to remove precipitated aggregates. Sample was purified using a HiTrapSP HF column (Cytiva) and eluted using a linear gradient from 50 mM to 1 M NaCl in 25 mM Na Phos pH 8.0, 0.5 mM TCEP. Fractions containing pure protein, as determined by SDS PAGE, were combined and dialyzed at 4°C in 25 mM Na Phos, 150 mM NaCl, 0.5 mM TCEP pH7.

### Isothermal Titration Calorimetry

Purified protein samples were dialyzed in 25 mM Na Phos, 150 mM NaCl, 0.5 mM TCEP, pH7 (dialysis buffer) at 4°C overnight. Proteins were 0.2*µm* filtered using 13 mm diameter PVDF filters (Thermo Fisher part no. CH2213-PV). Protein concentrations were determined from their ultraviolet absorbance and their calculated extinction coefficients [13]. NaCl was added to dialysis buffer, filtered, and used to bring experimental samples to 200 mM NaCl. Samples were left to equilibrate for twenty minutes before loading into a MicroCal VP-ITC. Experiments were performed at 25°C unless otherwise described, injection volumes were 8µL, and injection intervals were 5 minutes. Initial concentrations of proteins were as described throughout the paper, but typically 20*µM* protein was loaded in the syringe and 2*µM* of CSL was loaded in the calorimeter cell (with or without another protein in the cell in excess of 2*µM*). Raw thermogram injection peaks were baseline-corrected and integrated using NITPIC [19] and fit using the equations described in Appendix 1 or 3. Fitted parameters were from global fits of three experiments, and uncertainties are standard errors from the covariance matrix.

## Appendix 1. Solution of the cubic equation for CMA, and inclusion of the effects of dilution due to titration

As described above, when the mass-action expression for the ternary equilibrium constant is expressed in terms of total (rather than free) concentrations of CSL, MAML, and ANK (equation 5), the resulting expression is cubic in the concentration of the ternary complex. This is clearly revealed by rearranging equation 5 and collecting like terms:

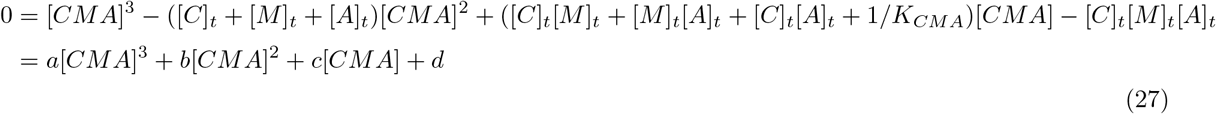

where

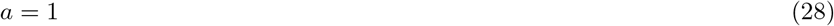

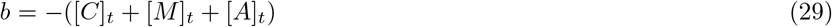

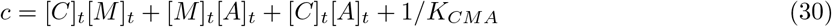

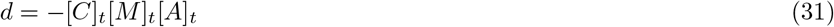

Although the three roots of equation 27 can be expressed analytically in terms of square and cube roots, these roots contain complex numbers and cannot be used for curve fitting. An alternative approach, which generates real roots, involves rearranging the cubic equation to eliminate the quadratic term. The roots of this rearranged (or “depressed”) cubic can be expressed using trigonometric functions rather than radicals. Obtaining a depressed cubic equation is achieved by a substitution of variables from [*CMA*] to *x*:

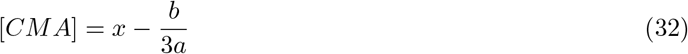

where *a* and *b* are as defined above. Substituting [CMA] in equation 27 with the right hand side of equation 32 and rearranging gives

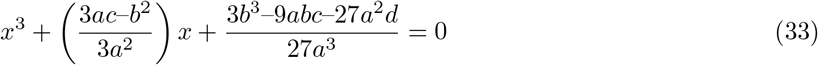

This is of the form

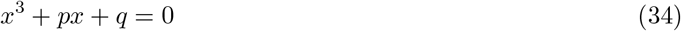

where

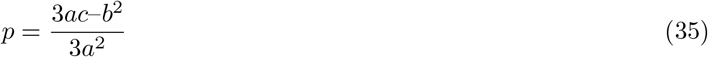

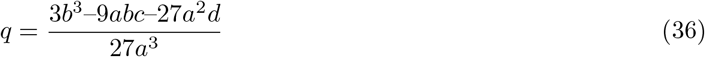

The three roots of the depressed cubic (equation 34) are given by the trigonometric formula

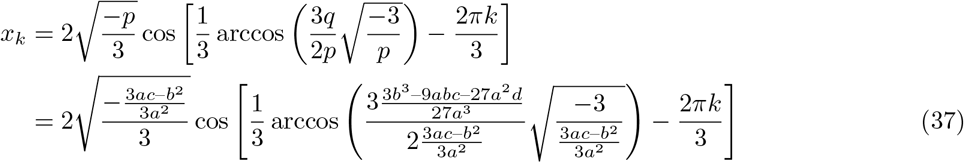

where *k* = 0, 1, or 2; for the ITC titrations here, the third root (*k* = 2) corresponds to positive values of [CMA]. Substituting *k* = 2 and *a* = 1 into equation 37 gives

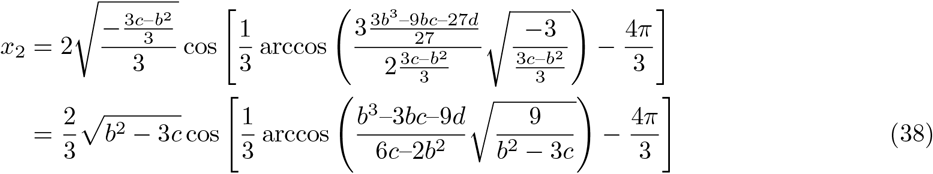

An expression for [*CMA*] can be obtained from *x*_2_ by substituting a rearranged version of equation 32, i.e.

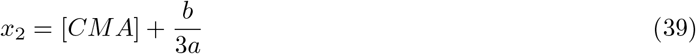

The resulting expression gives [CMA] as a function of the total concentrations [*C*]_*t*_, [*M*]_*t*_, and [*A*]_*t*_. As described in footnote 2, although total concentrations are uniquely determined from the known starting concentrations in the cell ([*C*]_0_, [*A*]_0_) and the syringe ([*M*]_0_), they need to be calculated after each injection to account for dilution and displacement effects. For proteins in the cell, each injection displaces some of the sample; the fraction of the sample displaced is equal to 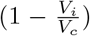 where *V*_*i*_ and *V*_*c*_ are the volume of the injection and cell, respectively. Cell proteins are diluted by this factor after each injection; thus, after injection *j*, the total protein concentrations are given by

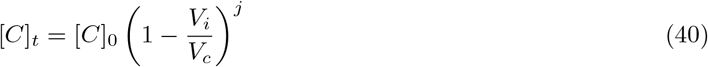

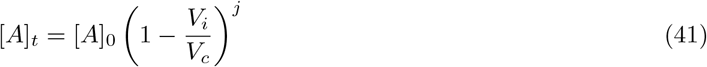

For the protein in the syringe (MAML in Figure 4), the first injection adds an amount of protein that is diluted by the ratio of the cell volume to the syringe volume, i.e.,

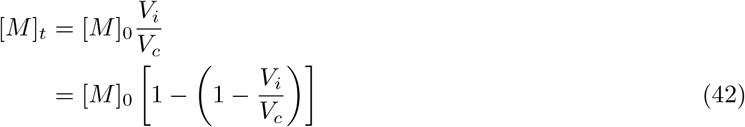

The rearrangement in the second line puts the [*M*]_*t*_ expression in the same form as expressions for subsequent injections. The second injection dilutes the MAML provided from the first injection (now in the cell) as in equations 40 and 41, and also adds another dilution from the syringe. Thus,

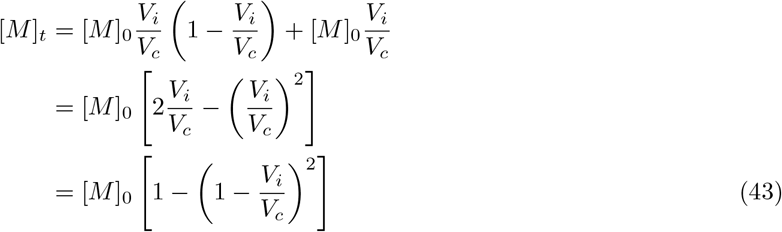

The third injection acts the same as the second, diluting MAML from the second injection and adding another dilution:

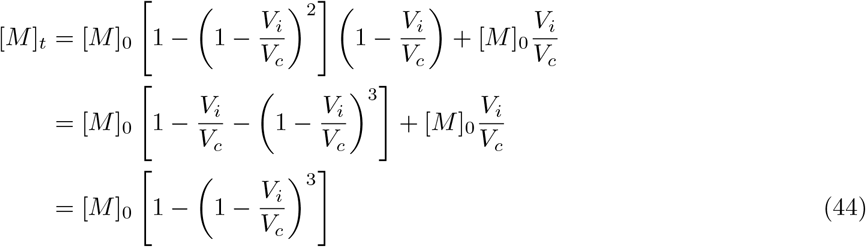

Comparing equations 42, 43, and 44 reveals a general formula for the total concentration of the syringe protein after the *j*^*th*^ injection:

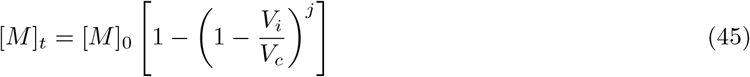

With these equations, an expression for the concentration of the ternary complex ([*CMA*]) can be generated with the following steps:

1. Substitute expressions for [*C*]_*t*_, [*M*]_*t*_, and [*A*]_*t*_ (equations 40, 45, and 41) into the equations for the coefficients to the cubic equation for [*CMA*] (*a, b, c*, and *d*, equations 28 - 31).
2. Substitute the modified cubic coefficients from step 1 into the equation for *x*_2_ (equation 38).
3. Substitute the modified expression for *x*_2_ from step 2 into the change-of-variable equation for [*CMA*] (equation 32).
4. Substitute the modified expression for [*CMA*] from step 3 into the heat equation (equation 3).

Although the resulting fitting equation (from step 4) is too long to fit on a page, it can easily be generated using the sympy module of python, and can be converted using the sympy.lambdify command to generate an equation that can be fit to ITC data (represented as a numpy array). Single ITC data sets are fit using the scipy.optimize.curvefit command. Multiple ITC data sets are globally fitted using the lmfit module. A jupyter notebook that generates this fitting function (along with sample data) is available on github.

## Appendix 2. Binding polynomials for RAMANK and MAML binding to CSL *in cis* and *in trans*

Analysis of ligand binding often requires expressions for fractions of macromolecules in different states of assembly. One convenient expression to represent such fractions is the binding polynomial [38, 2]. The binding polynomial is composed of statistical weights that represent the concentrations of ligation states of a macromolecule relative to its unbound state. Replacing concentrations with equilibrium constants generates a polynomial in powers of ligand concentration (or concentrations when there is more than one ligand).

Here we present binding polynomials that describe the saturation state of CSL with various ligands (RAM, ANK, RAMANK, and MAML). In these polynomials, unbound CSL serves as the reference state for the various bound forms of CSL. These polynomials are useful for calculating populations of CSL in various ligation states (Figures 8 and 9) and for generating expressions for numerical analysis of binding data. When RAM and ANK are *in trans* in the presence of MAML, the CSL binding polynomial has four terms:

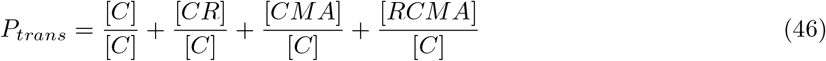

Substituting concentration ratios with equilibrium constants gives

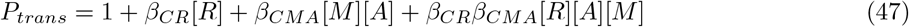

When RAM and ANK are *in cis* in the presence of MAML, the CSL binding polynomial has five terms:

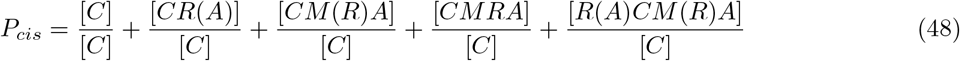

Substituting concentration ratios with equilibrium constants gives

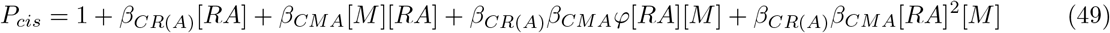

The fraction of total macromolecule that forms a particular species can be calculated by dividing the statistical weight of the species by the binding polynomial. For example, the fraction of CSL in that is in the bivalent ternary complex (CMRA) is

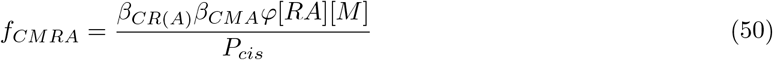

Likewise, the fraction of total CSL in the tetrameric complex R(A)CM(R)A is

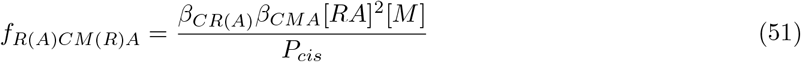

## Appendix 3. Numerical derivations for analysis of ITC data including the tetrameric R(A)CM(R)A complex

As described above, the fitted value of 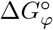 obtained from the titration of MAML into mixtures of CSL saturated with RAMANK indicates that at high concentrations of RAMANK, a non-neglibile amount of heterotetramer complex should form with two RAMANK, one CSL, and one MAML (R(A)CM(R)A, Figure 2), and should be included in fits of titrations of prebound CSL-RAMANK complexes with MAML. Modifying the equation for heats of injection (equation 3) to include this additional species gives

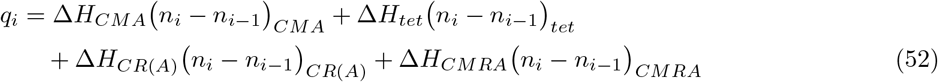

where the tetrameric species R(A)CM(R)A is abbreviated “tet”. The terms in parenthesis are the number of moles of each species in the cell at injection *i*. Substituting the mole number for each species with concentration gives

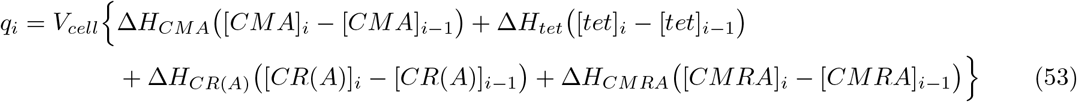

where *V*_*cell*_ is the volume of the calorimetry cell. Replacing molar concentrations of each species with species fractions gives

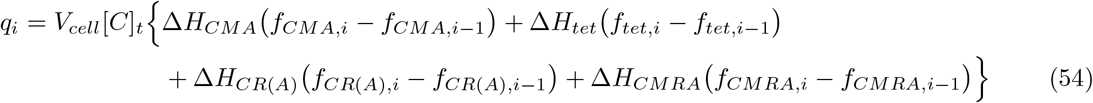

The species fractions can be calculated directly from the binding polynomial (Appendix 2), given a set of binding constants and unbound MAML and RAMANK concentrations. Unfortunately, calculating unbound concentrations from known total concentrations and equilibrium constants leads to an equation that is quartic in [*tet*] (and cubic in [*CMA*]). Rather than attempting to solve this equation analytically, we have used a numerical method to determine the concentrations of species needed to fit ITC titrations of MAML into CSL and RAMANK. This approach follows that of Vega *et al*. [35].

In the approach of Vega et al., the binding polynomial is used to generate expressions for concentrations of unbound species species in terms of total concentrations, and unbound concentrations are determined using a numerical root-finding algorithm. For titration of CSL and RAMANK with MAML, there are three such equations. For CSL,

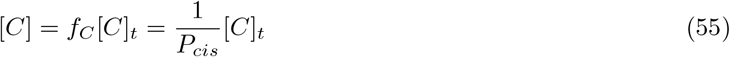

where *f*_*C*_ is the fraction of total CSL in the free state. In the expression on the right-hand-side, the one in the numerator is the statistical weight of free CSL, which is the reference state in the polynomial (see Appendix 2, equation 48).

An analogous expression for free MAML is derived from a conservation expression which states the total MAML concentration is equal to the sum of all species that include MAML,

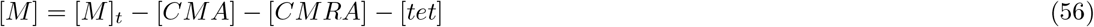

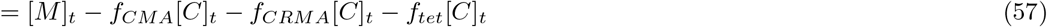

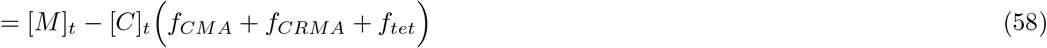

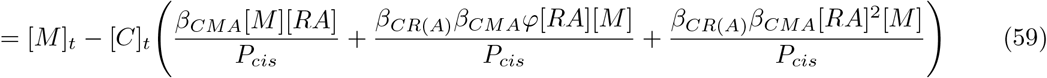

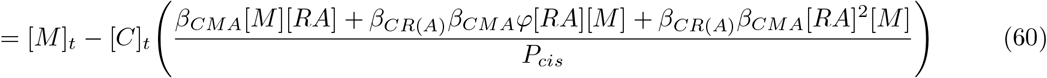

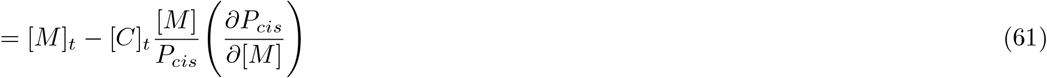

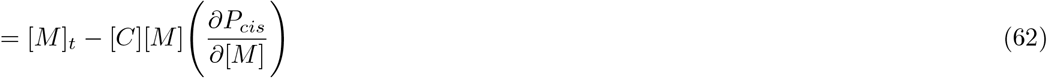

The reason this equation has more terms than that for [*C*] (equation 55) is that the binding polynomial *P*_*cis*_ is referenced to unbound CSL. The derivative in equations 61 and 62 provide a shorthand for the cluttered equations 59 and 65 [38, 2].

Likewise, the expression for free RAMANK is

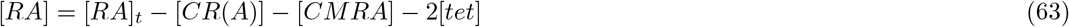

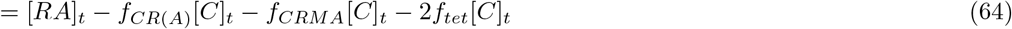

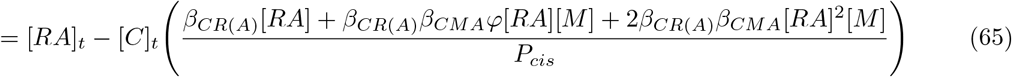

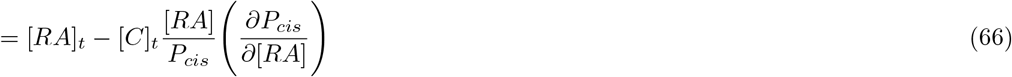

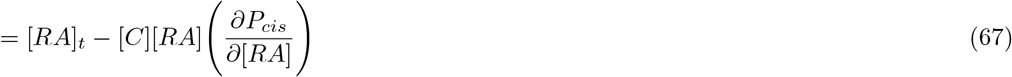

The factor of two multiplying [*tet*] accounts for the fact that there are two RAMANK molecules in the tetramer.

Rearranging the three expressions for unbound concentrations of CSL, MAML, and RAMANK (equations 55, 62, and 67) gives

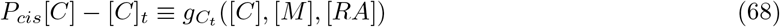

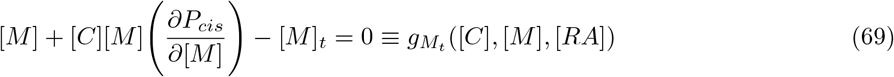

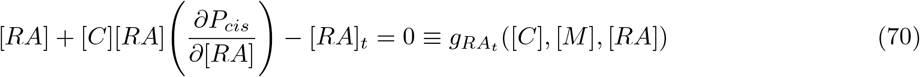

The *g* functions on the right-hand side is a shorthand to simplify the derivation below. Solving for [*C*], [*M*], and [*RA*] is equivalent to simultaneously finding the roots of equations 68, 69, and 70 given total concentrations [*C*]_*t*_, [*M*]_*t*_, and [*RA*]_*t*_. For a single-variable function *g*(*x*), the root can be found iteratively using the Newton-Raphson method. Starting with an initial guess *x*_0_ in the range of the true root *x*_*t*_, a better approximation to the root is given by

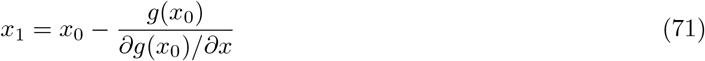

This method is repeated iteratively for some number of steps *k* at which a convergence criterion is satisfied:

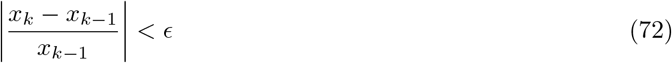

The same iterative approach can be used to simultaneously find the roots to equations 68, 69, and 70 where the initial guess *x*_0_ is replaced with a vector of initial guesses of concentration:

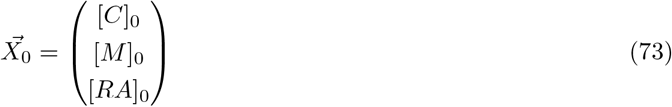

The function *g*_*x*_ is replaced by a vector 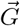 composed of the three functions:

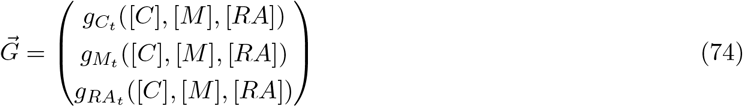

The derivative ∂*g*/∂*x* is replaced by the Jacobian matrix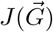:

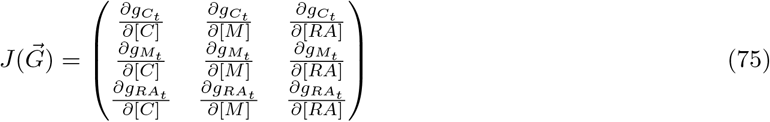

With these substitutions, the Newton-Raphson algorithm becomes

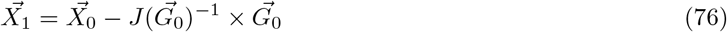

where 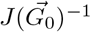 is the inverse of the Jacobian matrix evaluated at 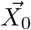. Estimates of [*C*], [*M*], and [*RA*] are iteratively updated using equation 76 taking the output of the previous iteration 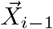 as input:

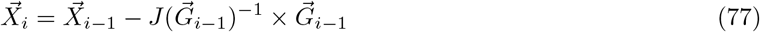

Optimization is terminated when three convergence criteria are simultaneously satisfied:

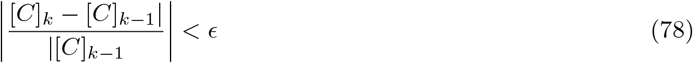

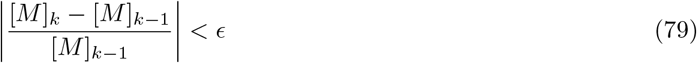

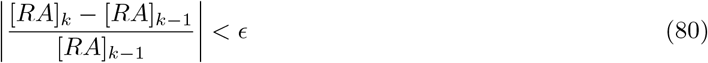

where *ϵ* is typically on the order of 10^−6^.

To fit ITC data, total concentrations ([*C*]_*t*_, [*M*]_*t*_, and [*RA*]_*t*_) are calculated for each injection using equations 40, 45, and 41 along with the starting cell and syringe concentrations ([*C*]_0_, [*M*]_0_, and [*RA*]_0_). These total concentrations are used in Newton-Raphson to find unbound concentrations [*C*], [*M*], and [*RA*] at each injection.

Combining these unbound concentrations with initial guesses for equilibrium constants^7^, the binding polynomial is evaluated at each injection, giving the species fractions needed to calculate heats of injection using equation 54. Calculated heats are subtracted from experimentally determined values and are supplied to the nonlinear least squares module lmfit [27] to optimize fit parameters (binding constants, enthalpies, and the incompetent fraction of MAML). With this approach, Newton-Raphson is rerun at every injection point in each round of nonlinear least squares optimization using updated fit parameters. Fitting single titrations, convergence times for lmfit are on the order of 80 seconds.^8^. A jupyter notebook containing the code for numerical fitting, along with some sample ITC data, can be downloaded from Github.

## Footnotes

1 Here Φ indicates hydrophobic residue

2 More precisely, the initial concentrations of the proteins in the cell ([*C*]_0_, [*A*]_0_) and syringe ([*M*]_0_) are known. Total concentrations after injection *i* are decreased from initial values by dilution and displacement associated with each injection. The conversion from initial concentrations to total concentrations is provided in Appendix 1.

3 Assuming 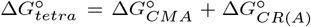, the estimate of 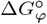 from 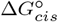 (Figure 5) can be used to calculate the relative amounts of bivalent ternary and tetrameric complexes (see Appendix 2). At 6*µM* free RAMANK and 3*µM* free MAML, CSL should form 77% bivalent ternary complex and 23% tetrameric complex.

4 For heterodimer formation, a free energy of −18.0 kcal/mol would result in saturation binding at micromolar concentrations in which heat peaks would drop abruptly from full amplitude to zero in a single injection.

5 Note that for this analysis, concentrations (i.e., [*M*] and [*A*]) are free (rather than total) concentrations

6 Though free MAML concentrations are not constant in actual ITC experiments

7 Or with updated values during the fit.

8 The slowest part of the numerical fitting procedure is calculating the Jacobian and its inverse, which only needs to be done once.

## Notes

### Competing Interest Statement

The authors have declared no competing interest.

### Summary of Updates

Some fits were updated, the abstract and title were changed for clarity, and text was added addressing the limits of detection for isothermal titration calorimetry.

