## Supplemental_Figures_and_Tables for "Bivalent interaction through an intrinsically disordered linker promotes transcription activation complex assembly in Notch signaling"

#### Supplementary Materials

Supplemental Figures S1-S4

Supplemental Tables S1-S2

#### Supplemental Figures

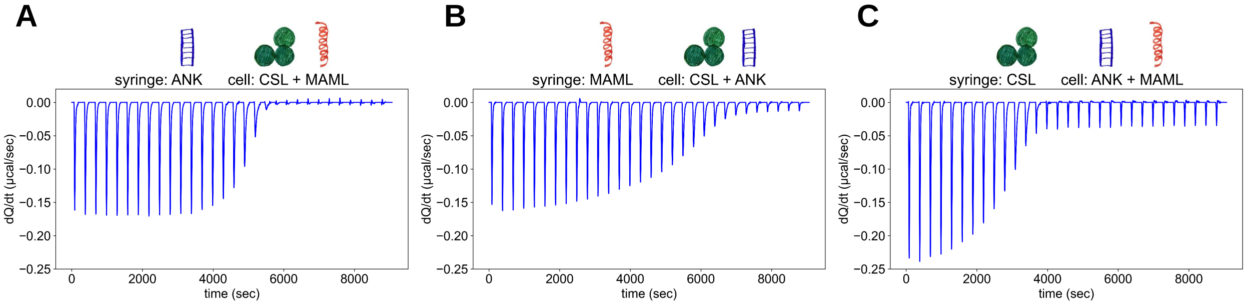


Figure S1: **ITC can detect formation of ternary complexes from ANK, MAML, and CSL.** (**A**) 2 $\mu M$ CSL and 35 $\mu M$ MAML titrated with 20 $\mu M$ ANK. (**B**) 2 $\mu M$ CSL and 8 $\mu M$ ANK titrated with 20 $\mu M$ MAML. (**C**) 2 $\mu M$ ANK and 8 $\mu M$ MAML titrated with 20 $\mu M$ CSL. Upper cartoons show positions of proteins in ITC experiment setup. Bottom panels show baseline corrected thermograms.

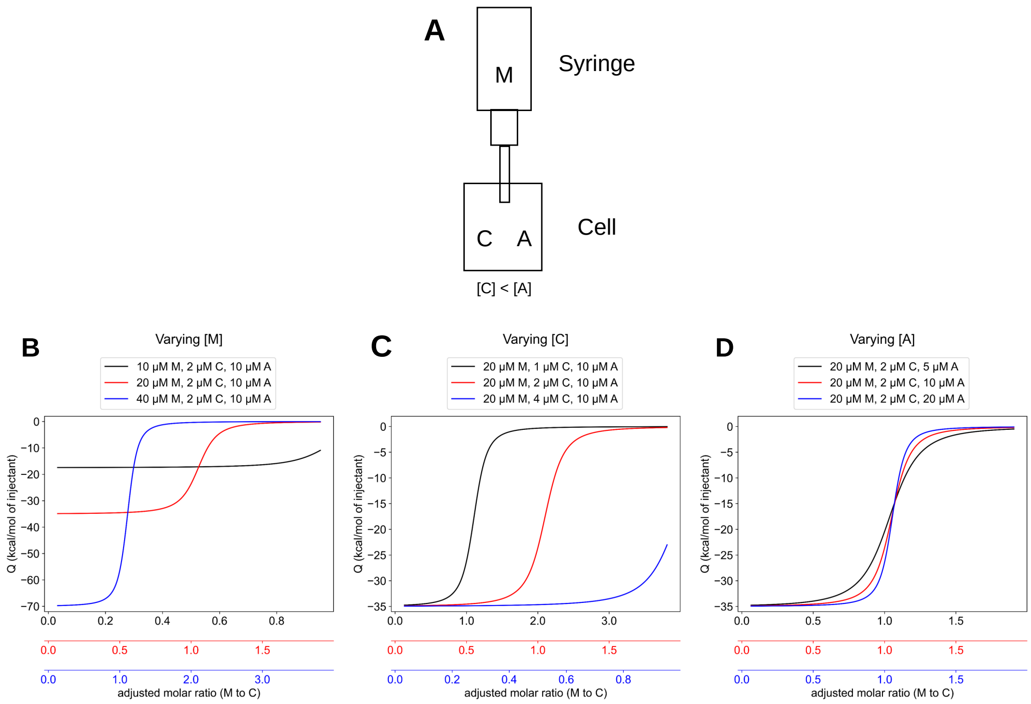


Figure S2: **The effects of variations in starting concentrations of reactants in obligate ternary complex formation on the profiles of integrated heat peaks.** (**A**) Schematic of an ITC experiment, with M in the syringe and C and A in the cell. In all simulations, C is substoichiometric to A. (**B**) Changing the initial concentration of the titrant ($M_0$) shifts both the y-intercept and the molar ratio of titrant to limiting cell ligand. (**C**) Changing the initial concentration of the limiting cell ligand ($C_0$) shifts molar ratio of titrant to limiting cell ligand, but does not affect the y-intercept. (**D**) Changing the initial concentration of the excess cell ligand ($A_0$) shifts neither the y-intercept nor the molar ratio of titrant to limiting cell ligand, but does shift the steepness of the integrated heat peaks' profile.

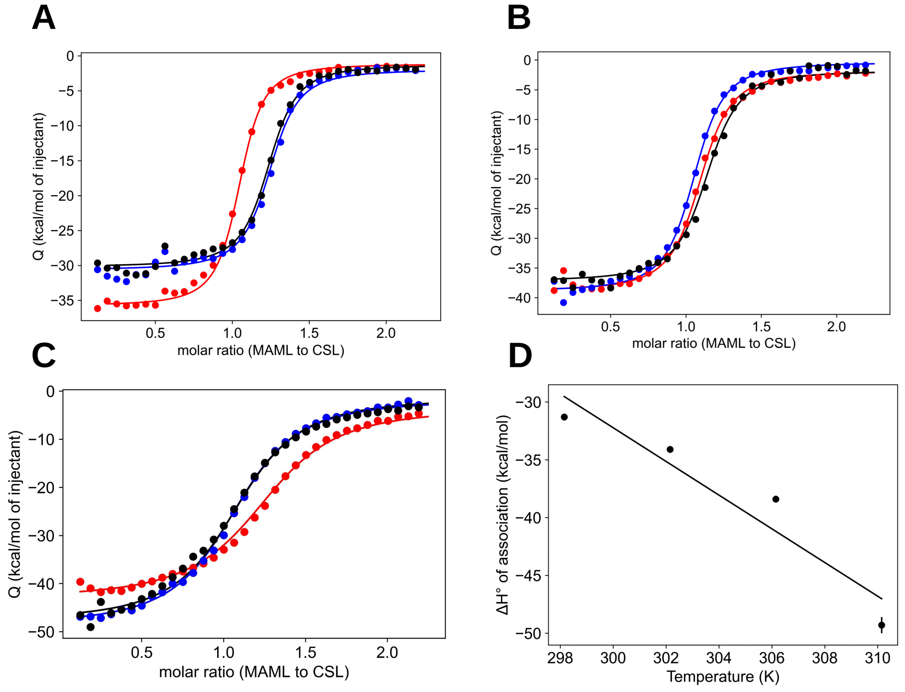


Figure S3: **Global fits of CSL-MAML-ANK at different temperatures to determine** $\Delta Cp_{CMA}$. (**A**) through (**C**) plot integrated heats of injection for three titrations at 29 °C, 33 °C, and 37 °C respectively where 20 $\mu M$ MAML in the syringe was titrated into 2 $\mu M$ CSL and 8 $\mu M$ ANK in the cell. Locally fitted competent fractions of MAML for (**A**) were 101% (red), 84% (blue), and 85% (black); for (**B**) were 96% (red), 100% (blue), and 92% (black); and for (**C**) were 80% (red), 95% (blue), and 94% (black). Solid lines are global fits using the obligate heterotrimer model (equation 3). (**D**) $\Delta H^{\circ}_{CMA}$ is plotted for 25 °C, 29 °C, 33 °C, and 37 °C. The slope of $\Delta H$ as a function of temperature gives ΔCp, the heat capacity change for CSL-ANK-MAML association (-1.46 kcal/mol K, ± 0.33).

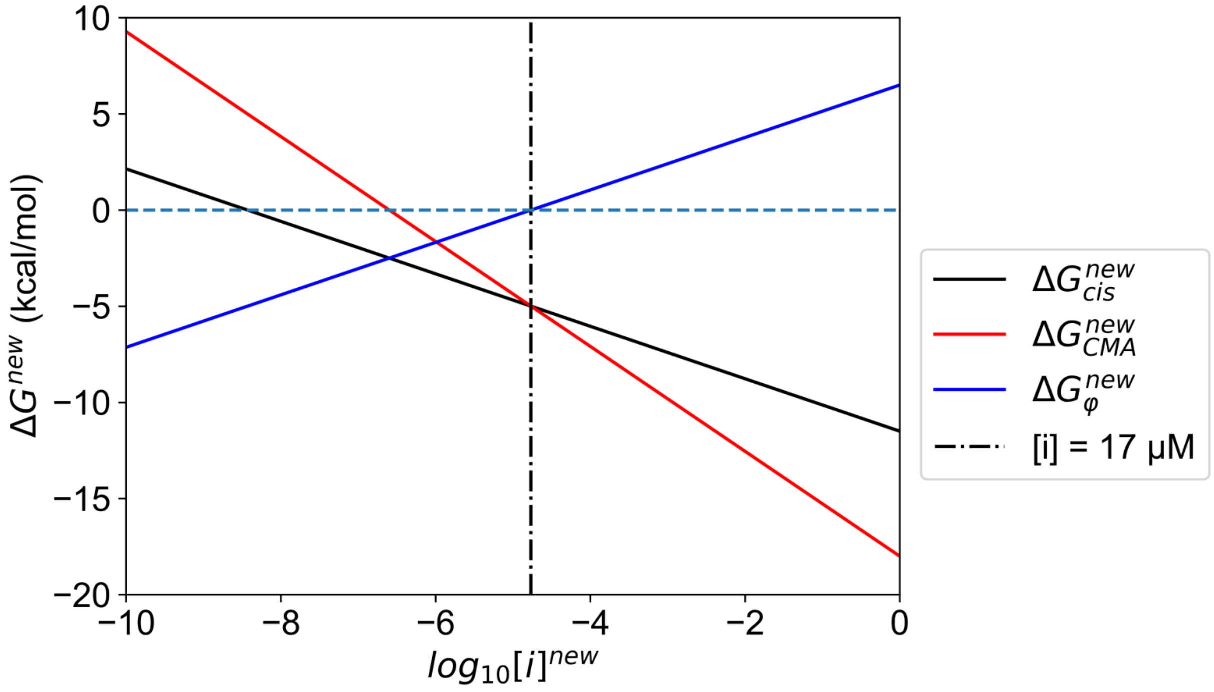


Figure S4: **Dependence of free energies of ternary complex formation on standard state concentrations.** $\Delta G^{new}_{cis}$, $\Delta G^{new}_{CMA}$ and $\Delta G^{new}_{\varphi}$ are calculated as a function of standard state concentration [i] using equations 24 - 26. Because $\Delta G^{new}_{cis}$ and $\Delta G^{new}_{CMA}$ describe bimolecular and trimolecular association respectively, they have different slopes and therefore intersect, intersecting when the standard state is equal to $C_{eff}$ (the vertical dash-dotted line). Below $C_{eff}$, more ternary complex is formed *in cis* than *in trans*, corresponding to a negative $\Delta G^{new}_{\varphi}$.

### Supplemental Tables

Table S1: **ITC experiments to dissect CSL:RAMANK:MAML assembly thermodynamics**

| # | Rxn | Syringe | Cell | $\Delta G^\circ$ | $\Delta H^\circ$ |
|---|---|---|---|---|---|
| 1 | $C + R \rightleftharpoons CR$ | RAM | CSL | $\Delta G^\circ_{CR} = -10.2 \pm 0.1$ | $\Delta H^\circ_{CR} = -20.1 \pm 0.3$ |
| 2 | $C + RA \rightleftharpoons CR(A)$ | RAMANK | CSL | $\Delta G^\circ_{CR(A)} = -10.5 \pm 0.1$ | $\Delta H^\circ_{CR(A)} = -19.0 \pm 0.2$ |
| 3 | $C + A \rightleftharpoons CA$ | ANK | CSL | n.d. | n.d. |
| 4 | $C + M \rightleftharpoons CM$ | MAML | CSL | n.d. | n.d. |
| 5 | $M + A \rightleftharpoons MA$ | MAML | ANK | n.d. | n.d. |
| 6 | $C + M + A \rightleftharpoons CMA$ | MAML | CSL + ANK | $\Delta G^\circ_{CMA} = -18.0 \pm 0.1$ | $\Delta H^\circ_{CMA} = -31.3 \pm 0.3$ |
| 7 | $CR + M + A \rightleftharpoons RCMA$ | MAML | CR + ANK | $\Delta G^\circ_{RCMA} = -18.3 \pm 0.1$ | $\Delta H^\circ_{RCMA} = -26.7 \pm 0.3$ |

Titrations were performed at 25 °C in 200 mM NaCl, 0.5 mM TCEP, 25 mM $Na_2HPO_4$, pH 7.0. For reactions 3-5, ITC thermograms show small heats that remain constant over the course of the titration, providing no evidence of binding. Units of $\Delta G^\circ$ and $\Delta H^\circ$ are kcal/mol. Errors are standard deviations on the mean from three technical replicates. .

Table S2: **ITC experiments to dissect energetics of bivalency**

| # | Rxn | Syringe | Cell | $\Delta G^\circ$ | $\Delta H^\circ$ |
|---|---|---|---|---|---|
| 8 | $CR(A) + M \rightleftharpoons CMRA$ | MAML | CR(A) | $\Delta G^\circ_{cis} = -11.6 \pm 0.1$ | $\Delta H^\circ_{cis} = -28.3 \pm 0.3$ |
| 9 | $CR(A) + M + RA \rightleftharpoons CMRA + R(A)CM(R)A$ | MAML | CR(A) | $\Delta G^\circ_{\varphi} = 6.5 \pm 0.2$ | $\Delta H^\circ_{\varphi} = 9.5 \pm 1.5$ |

Titrations were performed at 25 °C in 200 mM NaCl, 0.5 mM TCEP, 25 mM $Na_2HPO_4$, pH 7.0. Units of $\Delta G^\circ$ and $\Delta H^\circ$ are kcal/mol. Errors are standard error on the mean from at least three technical replicates. .
